# The heterogeneous selection landscape of genome evolution in prokaryotes

**DOI:** 10.1101/2025.11.26.690804

**Authors:** Roman Kogay, Svetlana Karamycheva, Nash D. Rochman, Yuri I. Wolf, Eugene V. Koonin

## Abstract

Evolution of prokaryote genomes appears to be defined by the interplay of selection for genome streamlining, deletion bias and selection for functional diversification. The previously observed overall positive correlation between the strength of selection, measured as the ratio of non-synonymous to synonymous nucleotide substitutions (*dN/dS*), points to diversification as the primary factor of prokaryote genome evolution. Here, we investigated the interplay between genome size and selection pressure by analyzing an expanded collection of closely related prokaryotic genomes, evaluating genome-wide selection by measuring *dN/dS* by using an accurate, phylogeny-based method and decomposing the resulting values into lineage-specific and gene-specific components. These analyses reveal a pronounced heterogeneity in the relationship between genome size and the strength of selection across the diversity of prokaryotes. Most bacteria display a positive correlation consistent with selection for diversification, whereas all analyzed archaeal lineages show strong negative correlation which is the signature of streamlining. These findings indicate that the selection regimes broadly vary across the diversity of prokaryotes rather than following a single, universal pattern. Genome streamlining, selection for functional diversity and drift in small populations are all important factors of evolution, their relative contributions depending on the population genetics and ecology of a given lineage.

## Introduction

Evolution of genome complexity that is reflected in the number of distinct genes is thought to be shaped primarily by the product of selection and effective population size (*Ne*) ^1^. Within this population-genetic framework and considering the universally observed excess of deletions over insertions (deletion bias) in the evolution of genomes ^2,3^, prokaryotic evolution is often viewed as being dominated by selection for genome streamlining ^1,4,5^. Indeed, some of the most abundant (very large *N_e_*) free-living bacteria, notably those adapted to oligotrophic marine conditions, such as members of the SAR11 clade, have the smallest genomes among free-living bacteria and evolve under strong purifying selection, in the streamlining regime ^4,6^. However, streamlining Is not the only major factor in the evolution of prokaryote genomes that exhibit broad diversity in gene number and composition, ranging over two orders of magnitude in size, from tiny, highly reduced genomes of endo- and ecto-symbionts ^7^, some of which encompass less than 200 genes, to gene-rich genomes of many free-living bacteria, some containing more than 11,000 genes ^8–11^.

With the sequencing of first genomes of obligate endosymbionts, one of the most striking observations was the pervasive reduction in their genome sizes ^12,13^. These drastically eroded genomes have lost a large fraction of metabolic and regulatory genes, retaining only core functions required for translation, replication and host dependency ^14^. Such extreme genome reduction appears to be a combined effect of genetic drift and a pronounced deletion bias ^3^. Small *N_e_*that is typical among bacteria with host-restricted lifestyles substantially limits the efficiency of selection, facilitating accumulation of deleterious deletions, which eventually results in a gradual genome contraction and functional simplification ^15^. Thus, genome contraction in prokaryotes can occur both under strong selection for streamlining and via a deletion ratchet under weak selection. In contrast, the selection for diversification favors the expansion of genetic and regulatory repertoire, promoting metabolic versatility, and regulatory complexity ^16^.

Generally, the strength of selection in prokaryotes is proxied by the ratio of non-synonymous to synonymous mutations (*dN/dS*) in protein-coding genes which reflects the strength of selection on the protein level ^17^, which decreases with increasing *N_e_* ^18^. Previous studies have demonstrated that *dN/dS* in prokaryotes generally tends to negatively correlate with the genome size (total number of genes) ^16,19–21^, being partially driven by underlying phylogeny ^22^, which appears to be poorly compatible with the streamlining regime of evolution. Therefore, it has been proposed and encapsulated in a mathematical model that positive selection for functional diversification that results in acquisition of multiple genes, primarily, via horizontal gene transfer (HGT), overrides both purifying selection for streamlining and the deletion ratchet in prokaryote evolution ^21^. However, due to the methodological limitations, these studies primarily estimated *dN/dS* from pairwise genome comparisons, summarizing the results with median or mean values. This approach obscures phylogenetic nonindependence within lineages and inadequately accounts for shallower and deeper divergences within the group. Furthermore, previous analyses were primarily limited to conserved housekeeping genes that can potentially misrepresent genome-wide selection landscape if core genes experience different constraints than accessory genes. Most importantly, averaging of the correlations due to limited genomic data obscured potential heterogeneity of the selection landscapes across the diversity of bacteria and archaea.

Here, we sought to investigate how genome size relates to selection pressure using an expanded collection of closely related prokaryotic genomes from the latest version of the Alignable Tight Genomic Cluster (ATGC) database, containing over 22,000 genomes ^23^. We evaluated genome-wide selection by measuring *dN/dS* for various orthologous gene families using a phylogeny-aware approach and partitioned those values into lineage-specific and gene-specific components. The results of these analyses reveal a notable heterogeneity in the relationship between genome size and the strength of selection across the diversity of prokaryotes. Strikingly, most bacterial lineages display a positive correlation consistent with selection for diversification, whereas all analyzed archaeal lineages show strong negative correlation that appears to be the signature of streamlining. We demonstrate that the observed trends persist after controlling for sampling bias and are robust across various functional categories of genes. These findings indicate that the selection regimes vary broadly across the diversity of prokaryotes rather than following a single, universal pattern. Genome streamlining, selection for functional diversity and drift in small populations are all important factors of evolution, their relative contributions depending on the population genetics and ecology of a given lineage.

## Results

### Lineage-specific and gene-specific selection in prokaryotes

Accurate estimation of *dN/dS* values requires alignments that include sufficient numbers of both synonymous and nonsynonymous mutations, while avoiding saturation that can obscure evolutionary rates ^24–26^. To achieve this balance, we initiated our analyses using the latest version of the ATGC database, in which each ATGC consisted of at least four genomes that shared ancestral gene synteny and stay below the saturation limit at synonymous sites. To further ensure a robust signal, we retained only those ATGC COGs that encompassed at least 32 mutations across the alignment and only those ATGCs where at least 70% of orthologs satisfied this condition. Under these criteria, the final dataset consisted of 791 ATGCs (**Figure S1**), including 19 archaeal and 772 bacterial ones. Genome sizes within this collection of ATGCs spanned more than two orders of magnitude: the smallest genomes belonged to ATGC1045 containing members of the genus *Nasuia* (median length of 0.11 Mb), whereas the largest genomes were found among *Streptomyces* species in ATGC0548 (median length of 12.02 Mb) (**Table S1)**.

To validate the reliability and accuracy of *dN/dS* estimations for gene alignments using IQ-Tree ^27^, we conducted simulations that mimicked empirical patterns of sequence evolution in representative bacterial lineages. Specifically, we used the phylogeny of ATGC0001 (comprising *Escherichia*, *Salmonella* and *Citrobacter*) as a guide tree and generated 2,724 artificial codon alignments. For each simulated alignment, we used the observed substitution patterns (*dN/dS* and transition/transversion ratio), alignment length, and branch lengths estimated from the corresponding clusters of orthologous genes (COGs) ^28^. The resulting simulated alignments were then reanalyzed with IQ-Tree using the same mode selection procedure as applied to empirical alignments. The results demonstrate that IQ-Tree accurately recovered both *dN/dS* (Spearman’s R = 0.96, asymptotic p-value < 1 ∗ 10^−16^; **Figure S2A**) and transition/transversion ratios (Spearman’s R = 0.90, asymptotic p-value < 1 ∗ 10^−16^; **Figure S2B**), confirming its suitability for large-scale analyses of the selection regime in the evolution of bacterial and archaeal genomes.

To estimate the genome-wide selective constraints across the entire set of genes, including those that are not represented in (nearly) all ATGCs and to characterize the selection regimes more thoroughly, we decomposed *dN/dS* values across all COGs into components reflecting lineage (ATGC)-specific constraints, gene (COG)-specific constraints, and residuals. First, each ATGC COG was linked to the prokaryote-wide COGs^28^ (pCOGs here) by sequence similarity; when several paralogous ATGC COGs mapped to the same pCOG, the ATGC COG with the largest number of genes was chosen as the representative. For each 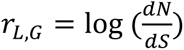, where *L* corresponds to lineage (ATGC), and *G* corresponds to gene (pCOG), we modeled the data as:

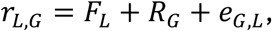

where *F_L_* is the lineage-specific component, *R_G_* is the gene-specific component, and *e*_*G*,*L*_ is the residual term (since *r*_*L*,*G*_ is in log scale, *F_L_* and *R_G_* are logarithms of the multiplicative factors). The parameters *F_L_* and *R_G_* were estimated iteratively by minimizing the sum of squared residuals:

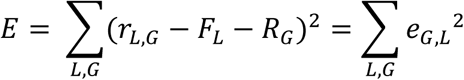

At each iteration, the mean of the residuals was calculated across all COGs for each ATGC to update *F_L_*, and then, across all ATGCs for each COG to update *R_G_*. These steps were repeated until convergence (change in *E* between iterations lower than 10^−6^) or a maximum of 1,000 iterations.

When estimating the strength of selection, it is generally assumed that selective forces act relatively uniformly across the evolutionary unit under study. To assess whether this assumption held for the ATGCs, we initially identified 30 ATGCs for which the phylogenetic tree could be partitioned into two subtrees, each containing at least 20 genomes with sufficient phylogenetic depth (see Methods). For each of these ‘splittable’ ATGCs, we calculated the *dN/dS* values for all COGs across the ATGC as a whole and separately for each of the two subtrees. In some ATGCs, selection patterns appeared visually uniform between the two subtrees, whereas others exhibited notable differences (**Figure S3**). To test whether these differences could arise from stochastic variation, we performed a bootstrap analysis. For each splittable ATGC, we randomly selected 100 COGs and calculated *dN/dS* values for 100 bootstrap replicates, measuring the proportion of samples that exceeded the value observed for the corresponding COG in the full ATGC (**Figure S4**). Although some variation in selection between subtrees was evident, in most ATGCs, only a small fraction of genes showed significant interclade differences within ATGCs.

The estimated ATGC-specific (lineage-specific) constraints suggest that only 6 of the 30 splittable ATGCs display moderate to strong heterogeneity of the selection pressures (>40% difference between subtrees; **Table S2**). Thus, most ATGCs exhibited relatively uniform selective regimes, validating their suitability as coherent units for comparative analyses of selection.

### Different selection regimes across the diversity of prokaryotes

To quantify the relationship between selection strength and genome size in bacteria and archaea, we aggregated *dN/dS* values from all COGs for 791 retained ATGCs, computed their ATGC-specific selective constraints, and used each ATGC’s median genome size as a representative (**Figure S5**). In this analysis, the ATGC-specific constraint was used as the proxy for selection strength in order to isolate the signal of potentially different selection regimes across bacterial and archaeal lineages. For the entire set of the ATGCs, a significant positive correlation between the genome size and the ATGC-specific constraint was observed, in agreement with previous studies ^16,19–21^ and consistent with selection for diversification (Spearman’s R = 0.26, 95% CI = 0.19 – 0.34; asymptotic p-value = 5 ∗ 10^−14^) (**Figure 1A**). Furthermore, also in agreement with the previous study ^22^, the ATGCs with intermediate genome sizes showed a wide spread in selective constraint values, whereas the largest genomes consistently exhibited moderate constraint.

**Figure 1.**
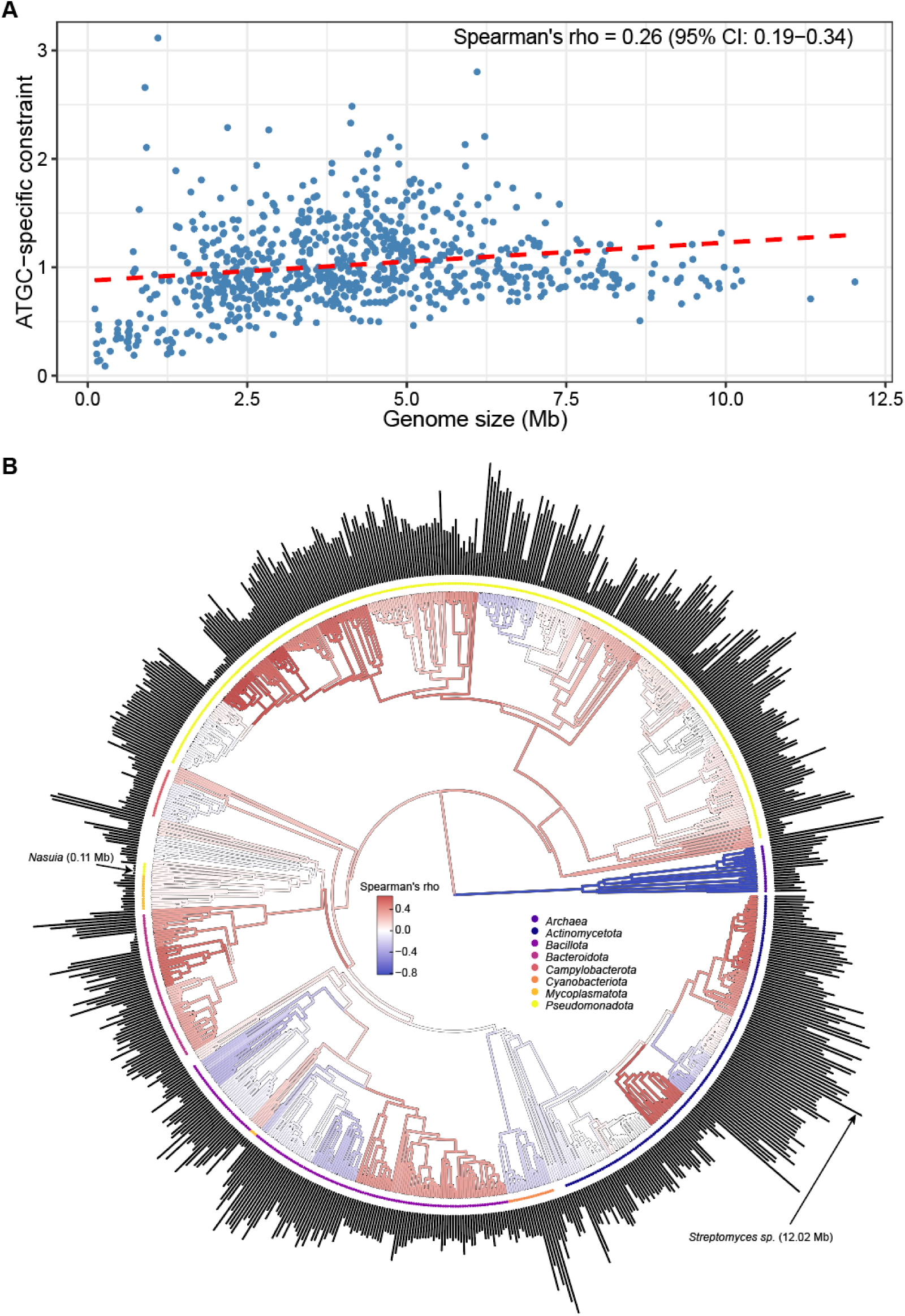
Relationship between genome size and ATGC-specific constraints across prokaryotes. **(A)** Scatterplot of genome size versus ATGC-specific constraint, which represents lineage-level selective pressure estimated from the decomposition of *dN/dS* values across orthologous genes. Each point corresponds to a bacterial or archaeal ATGC, with the genome size represented by the median value for that ATGC. **(B)** Distribution of Spearman’s correlation coefficients between genome size and ATGC-specific constraint across the prokaryotic phylogeny. Major bacterial phyla containing more than 15 ATGCs, as well as Archaea, are highlighted with distinct colors. Projecting bars indicate median genome sizes for individual ATGCs. Internal nodes are colored according to the correlation among their descendant lineages; only nodes with more than 15 descendants are directly colored, whereas others inherit the color of their parent node.

Mapping the Spearman’s R values between genome size and ATGC-specific selective constraint onto the phylogeny revealed pronounced heterogeneity across the prokaryotic tree, with the most pronounced contrast observed between the two prokaryotic domains of life, Bacteria and Archaea (**Figure 1B**). Bacteria, which dominate the ATGC dataset, showed a significant positive correlation between genome size and selective constraints (Spearman’s R = 0.28, 95% CI = 0.20 – 0.35; asymptotic p-value = 6 ∗ 10^−15^) (**Figure 2A**), implying selection for diversification. In contrast, *Archaea* exhibited a strong negative correlation between genome size and ATGC-specific selective constraints (Spearman’s R = -0.82, 95% CI: -0.93 – -0.64; asymptotic p-value = 1.8 ∗ 10^−5^) (**Figure 2B**). Similar to a previous report showing a positive correlation between *dN/dS* and genome size in *Archaea* ^29^, our results support the notion that larger archaeal genomes evolve under weaker purifying selection, which is consistent with the streamlining selection regime. To test whether the strong negative correlation between genome size and selection strength observed for *Archaea* could reflect sampling bias and imbalance in the analyzed datasets, we resampled bacterial ATGCs 10,000 times (n=19) and calculated the correlation coefficient for each draw. None of the samples had a R value less than -0.82, with the minimum observed value of -0.69, indicating that the negative correlation between genome size and the strength of selection in archaea could not be explained by sampling effects (**Figure S6**).

**Figure 2.**
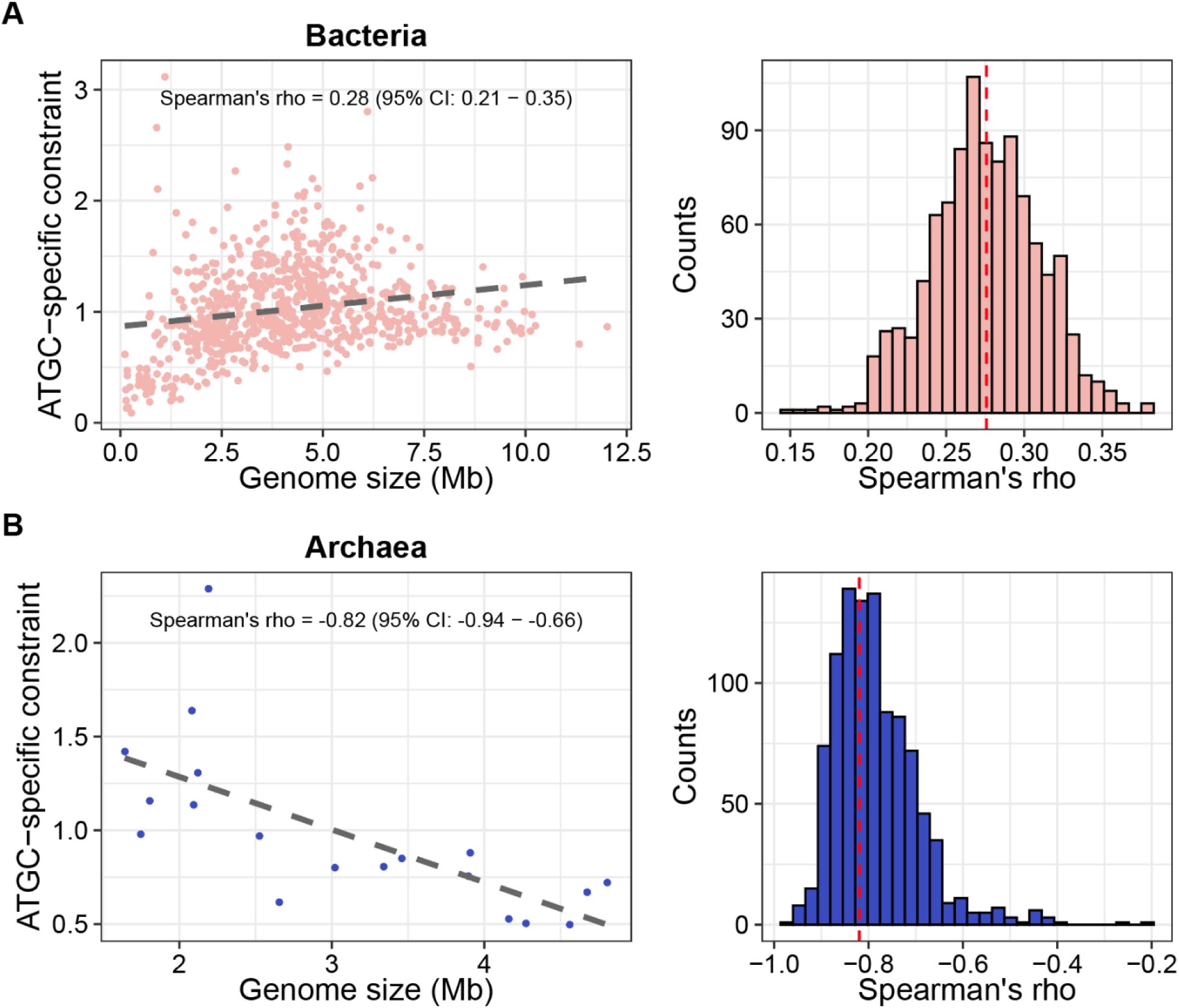
Contrasting relationships between genome size and ATGC-specific constraints in Bacteria (A) and Archaea (B). Left panels show scatterplots illustrating the relationship between ATGC-specific constraint and genome size. Right panels display the distributions of Spearman’s correlation coefficients obtained from 1,000 bootstrap replicates, which were used to calculate confidence intervals.

Although the overall correlation between genome size and selection strength among bacteria supports the selection for diversification regime, a more granular examination reveals pronounced heterogeneity across bacterial taxa. At the phylum level, strong positive correlation was observed for the vast phylum *Pseudomonadota* (Spearman’s R = 0.38, 95% CI = 0.27–0.49; asymptotic p-value = 1.1 ∗ 10^−12^ and for *Bacteroidota* (Spearman’s R = 0.35, 95% CI = 0.09-0.59; asymptotic p-value = 0.006), whereas for the other phyla, no significant relationship was detected (**Table S3)**. However, a finer taxonomic resolution reveals additional pockets of the diversification regime, such as orders *Mycobacteriales* (Spearman’s R = 0.41, 95% CI = 0.09 – 0.66; asymptotic p-value = 0.007) and *Lactobacillales* (Spearman’s R = 0.31, 95% CI = -0.01 – 0.56; asymptotic p-value = 0.031); these signals become masked when aggregated with other clades from the same phyla (**Figure 1B**, **Table S3**). Negative correlation implying streamlining is rare among the analyzed *Bacteria*, prominently observed only in the order *Bacillales* (Spearman’s R = -0.33, 95% CI = -0.52 – 0.12; asymptotic p-value = 0.009).

### Consistency of selection regimes across functional groups of prokaryotic genes

We further tested whether selection regimes varied among genes with different functions by grouping COGs into four classes: i) information storage and processing, ii) cellular processes and signaling, iii) metabolism, and iv) poorly characterized genes, and computing ATGC-specific selective constraints for each class independently. All four classes showed comparable overall positive correlations with genome size (Spearman’s R = 0.23 – 0.28, asymptotic p-values < 2.7 ∗ 10^−11^) (**Figure 3**), and near perfect concordance with the genome-wide ATGC-specific constraints (Spearman’s R = 0.97 – 0.99, asymptotic p-values < 1 ∗ 10^−16^) (**Figure S7**). Thus, selective pressures appear to be broadly shared across functional classes of genes. At finer resolution, constraints estimated for individual COG categories also closely matched genome-wide constraints and one another, reinforcing the conclusion of a uniform selective regime for most genes (**Figure S8**). A notable exception is the X category (mobilome), which showed a much weaker correlation with other COG categories (**Figure S8**). This deviation suggests that mobilome genes are substantially affected by other evolutionary forces, such as spread via HGT, making them less tightly coupled to the genome-wide selection landscape.

**Figure 3.**
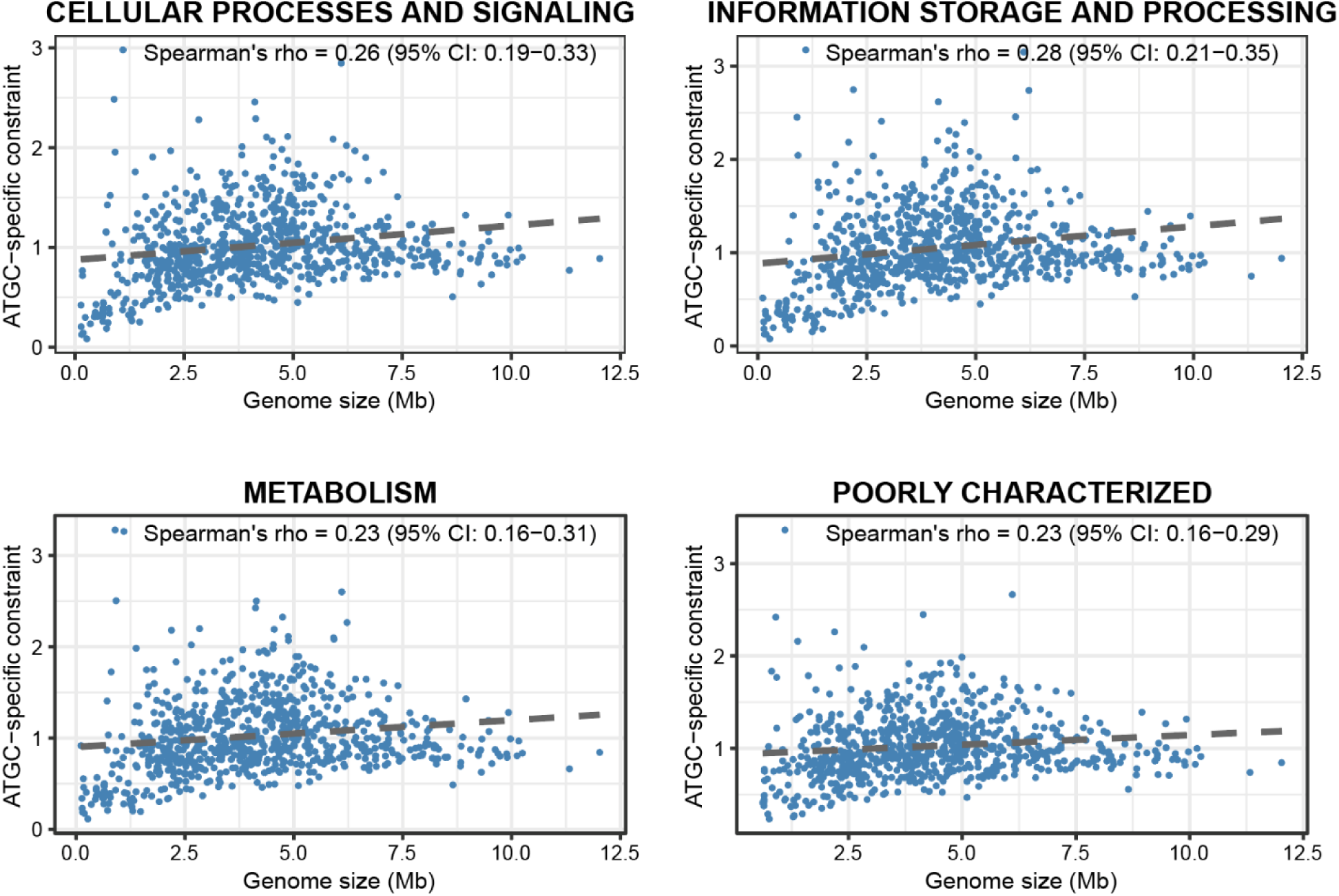
Relationship between genome size and ATGC-specific constraints across major functional categories of genes (COGs). ATGC-specific constraint represents lineage-level selective pressure estimated from the decomposition of *dN/dS* values within each functional subset. Only ATGCs with *dN/dS* estimates for at least 10 COGs were included in the analysis.

### Association between gene flux and selective constraints

To evaluate the association between the selection regimes and the overall genome dynamics, we quantified the relationship between gene flux rates and ATGC-specific selective constraints. Specifically, we tested whether the rates of gene gain, gene loss, and overall flux (gain/loss ratio) correlated with the strength of selection across ATGCs. No significant correlation was identified between the selective constraints and gene gain (Spearman’s R= 0.01, 95% CI: -0.06 – 0.09; asymptotic p-value = 0.78) or gene loss rate (Spearman’s R = -0.02, 95% CI: -0.10 – 0.05; asymptotic p-value = 0.52). In contrast, a significant positive correlation was observed between the selective constraints and the overall gene flux rate (Spearman’s R = 0.21, 95% CI: 0.14 – 0.27; asymptotic p-value = 3.3 ∗ 10^−9^) (**Figure 4A**). Thus, genomes that experience intensive gene flux are subject to significantly stronger selective constraints than more stable genomes. Notably, the gene flux itself did not significantly correlate with the genome size (Spearman’s R = -0.05, 95% CI: -0.12 – 0.02; asymptotic p-value = 0.15), indicating that differences in genome sizes between ATGCs are not determined by a simple imbalance between gene acquisition and loss. When mapped to the prokaryote phylogeny, the correlation between the selective constraint and flux appeared to be more uniformly distributed across clades than the correlation between constraint and genome size (**Figure 4B**).

**Figure 4.**
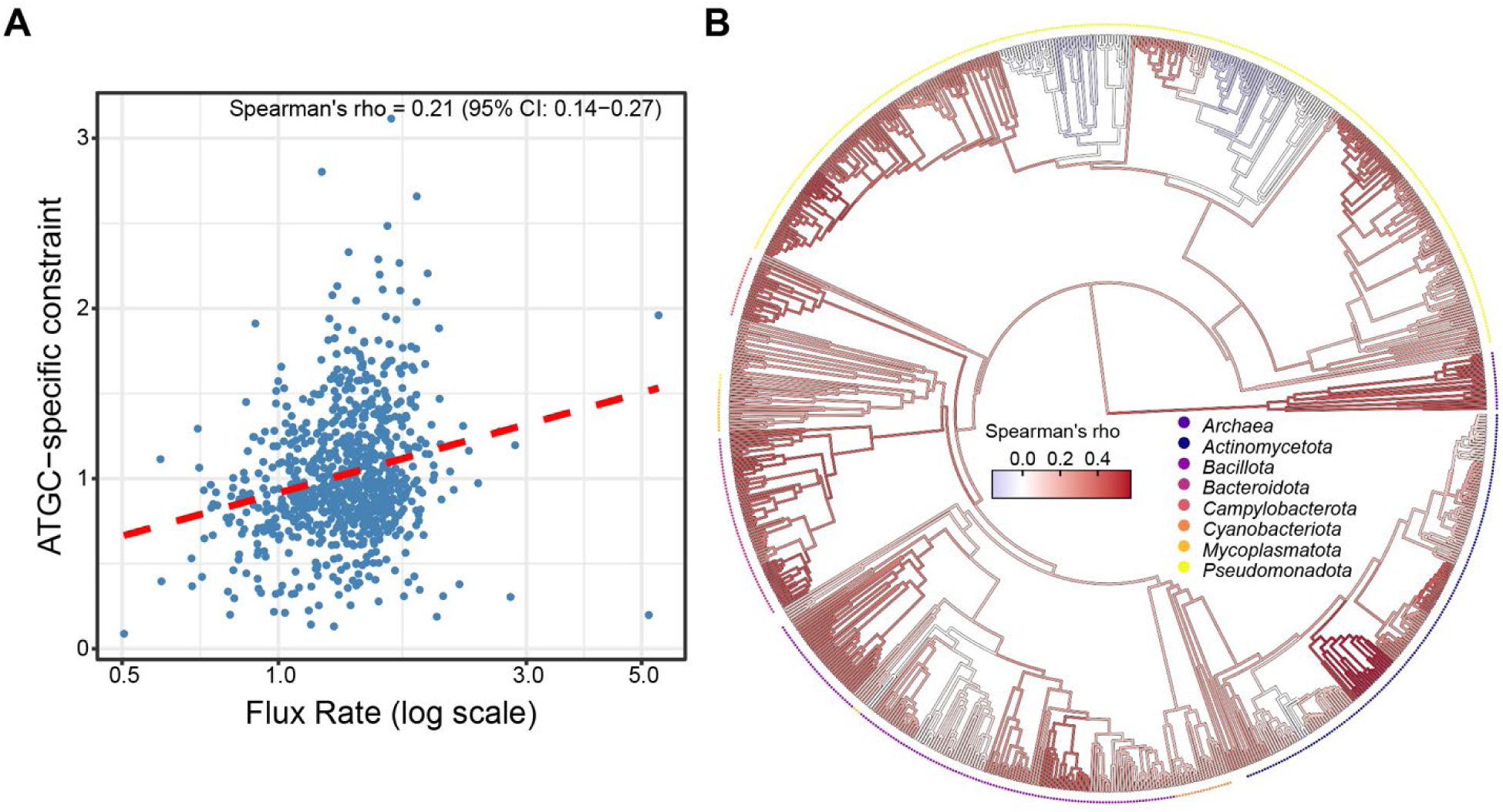
Relationship between gene flux rate and ATGC-specific constraints across prokaryotes. **(A)** Scatterplot of gene flux rate (log scale) versus ATGC-specific constraint, where each point corresponds to a bacterial or archaeal ATGC **(B)** Distribution of Spearman’s correlation coefficients between flux rate and ATGC-specific constraint across the prokaryotic phylogeny. Major bacterial phyla containing more than 15 ATGCs, as well as Archaea, are highlighted with distinct colors. Internal nodes are colored according to the correlation among their descendant lineages; only nodes with more than 15 descendants are directly colored, whereas others inherit the color of their parent node.

## Discussion

Previously, the evolution of prokaryote genomes was primarily considered to be driven either by selection for genome streamlining in large populations, as dictated by straightforward population genetics theory, or by selection for functional diversification as suggested by the observations on the positive correlation between selection strength and genomes size ^1,5,16,19–22^. These aggregate approaches were necessitated in part by the lack of a sufficient number of groups of closely related genomes, for which selection could be accurately estimated, and in part by the relatively crude methodology that estimated selection from pairwise genome comparisons. Here we extended and refined the analysis of genome-wide selection regimes in prokaryotes by exploring ATGCs representing a broad diversity of bacteria and archaea, applying a more accurate, phylogenetic approach for *dN/dS* estimation, and decomposing selective constraints into gene-specific and lineage-specific components.

This analysis validated the previous bulk estimates but showed that such estimates have been masking at least three distinct regimes of prokaryote evolution. The most striking observation is the dichotomy between the evolutionary regimes of the two domains of prokaryotes, bacteria and archaea. Whereas most bacterial taxa showed a positive correlation between selective constraints and genome size that implies selection for diversification, defining the overall correlation value, in archaea, a strong negative correlation was observed, suggestive of genome streamlining as the dominant selective factor. The observed negative association between selection constraint and genome size in *Archaea* is in an agreement with a previous report ^29^, and reinforces the idea that archaeal genome evolution follows a distinct selective regime compared to *Bacteria*. Although there are many fewer archaeal ATGCs than bacterial ones, the difference between the selection regimes cannot be explained by a sampling error because extensive resampling of bacterial data did not reveal any subsets with nearly as strong streamlining signal as that observed for archaea. However, apart from the pronounced differences between bacteria and archaea, substantial heterogeneity was observed within the bacterial domain, with some groups showing no significant dependency of selection on genome size, and one order exhibiting the signature of streamlining.

The causes of the pronounced dichotomy between the selective regimes of *Archaea* and *Bacteria*, and more generally, the heterogeneity of the selection landscapes among prokaryotes, remain to be explored and elucidated. Because selection constraints are inferred from the respective *dN/dS* values, they potentially can be driven by multiple underlying factors including effective population size, HGT rate, environmental stability and dynamics, and metabolic adaptation ^30,31^. Thus, a plausible explanation of the heterogeneity of the evolutionary landscape could be that, for generalists, primarily, bacteria, that occupy ecologically complex niches and have access to large gene pools and diverse interaction partners, the defining selective factor is functional diversification via acquisition of multiple genes through HGT ^32,33^. In contrast, for specialists, such as most archaea and some groups of bacteria, that typically inhabit stable, resource-limited or extreme environments, where high *N_e_* and limited ecological opportunities reduce the benefits of genetic innovation, streamlining appears to be the optimal evolutionary strategy ^34,35^. The interplay between environmental pressures and population genetics parameters likely amplifies differences in selection regimes over long timescales.

Compatible with the generalists vs specialists perspective, we observed a significant positive correlation between ATGC-specific selective constraints and the gene flux rate, but not with gene gain rate or loss rate separately, indicating that selection operates on the overall balance of opposing genome dynamics events, and is particularly strong in organisms with highly dynamic genomes. Gene acquisitions and deletions likely operate in a dynamic equilibrium that is shaped by both adaptive and neutral processes ^31,36^. Consequently, for lineages that evolve under selection for diversification, intense gene flux provides a constant source of potentially beneficial genes, some of which are then integrated into molecular networks and start evolving under purifying selection.

Although the positive correlation between genome size and selective constraints that is observed in most bacteria implies that many accessory genes are functionally relevant and evolve under strong purifying selection, this relationship is non-monotonic. Beyond an upper-intermediate genome size, the increase in selective constraint plateaus and can even decline (**Figure 1A**, **Figure 3**), suggesting limits to the adaptive value of an expanding gene repertoire. This nonlinearity implies diminishing returns for adding genes after a certain point, due to factors such as functional redundancy, increased metabolic burden and other costs that selection cannot offset, leading to weaker constraint ^37^. Furthermore, integration of new genes into the increasingly complex regulatory and metabolic networks require substantial molecular coordination, which may not be feasible beyond a certain genome size ^38,39^, making unlikely the fixation of acquired genes.

The lineage-specific selective constraints are highly consistent across functional classes of genes, scaling similarly with genome size. This covariation implies that, despite the broad variance of gene evolution rates, in most prokaryotes, changes in selection strength occur predominantly at a system-wide level, in accord with the concept of the universal pacemaker of genome evolution ^40^. The genes associated with the mobilome are the only prominent exception, showing a substantially weaker correlation with genome-wide constraints. This decoupling of the mobilome from core cellular processes reflects the selfishness of mobile elements that propagate relying on their own molecular mechanisms rather than directly contributing to the overall fitness of prokaryotes ^41^.

Overall, our present work shows that genome size evolution in prokaryotes reflects a continuum of lineage-specific selective regimes rather than a single, universal evolutionary trend. The interplay between ecological conditions, population genetics, and molecular mechanisms can result in diverse evolutionary trajectories, highlighting the complexity and adaptability of prokaryotes. However, more theoretical work on models of evolution along with empirical analysis of genome evolution in diverse groups of bacteria and archaea are required to elucidate the underlying mechanisms of evolutionary processes that shape the composition of prokaryote genomes.

## Methods

### Genomic dataset and *dN/dS* calculation

A total of 929 ATGCs were extracted from the ATGC database. Each genome was annotated by performing a competitive PSI-BLAST search of COG profiles ^28^. When multiple orthologous groups within the same ATGCs mapped to the COG, the group containing the largest number of sequences was retained. The number of individual mutations in each COG was estimated by first reconstructing a phylogenetic tree from codon alignments using FastTree (gamma + GTR model) ^42,43^, and then multiplying the total tree length by number of sites in corresponding alignment. Only alignments with at least 32 mutations were retained for downstream analyses.

*dN/dS* values were calculated using the IQ-Tree with ModelFinder ^27,44^, restricting search to models incorporating gamma distribution to account for variability in substitution rates among sites. A total of 2,724 alignments were simulated using AliSim ^45^, employing empirical molecular values from 2,724 COGs of ATGC0001, including their observed *dN/dS* and transition/transversion ratios. Simulations were based on the phylogenetic tree of ATGC0001 that was pruned to match the taxa present in empirical sequences with 5,000 codons per alignment. Subsequently, 5,000 codons were trimmed to match the length of the corresponding real alignments.

### Identification of splittable ATGCs

The phylogenetic tree of each ATGC was examined to identify branches that could split the tree into two subtrees, each with a phylogenetic depth greater than 0.2 and containing at least 20 genomes. If multiple branches met these criteria, the one that maximized the number of genomes on both subtrees was selected. During the analyses of 32 retained ATGCs, subtrees from 2 ATGCs produced largely unreliable *dN/dS* values (more than 30% equal to 1) and were excluded, leaving 30 ATGCs for subsequent analyses.

### Decomposition of *dN/dS* values into lineage-specific and gene-specific components and correlation analyses

Lineage- and gene-specific components of *dN/dS* were estimated using an iterative decomposition approach. For a COG *G* in ATGC *L*, the model for its observed log-transformed *dN/dS* ratio (*r*_*G*,*L*_):

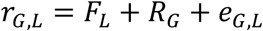

where *F_L_* and *R_G_* are lineage- and COG-specific constraints respectively, and *e*_*G*,*L*_ are residuals. Constraints were initialized at zero and were updated iteratively until convergence of the sum of squared residuals. Outliers with unlikely residuals under a normal approximation were removed, and the decomposition was repeated to obtain final estimates of constraints and residuals. Confidence intervals between ATGC-specific constraints and other metrics were estimated by performing 1,000 bootstrap replicates, followed by Bias-Corrected and Accelerated bootstrap (BCa), to account for bootstraps distribution and skewness ^46^.

### Estimation of gene flux rate

The data on the representation of ATGC genomes in COGs were converted into presence-absence binary sequences in FASTA format. Rooted ATGC trees together with the binary FASTA files were used as input for locally installed GLOOME software ^47^. Reconstructed ancestral states were mapped onto the corresponding phylogenetic tree. For each COG, the tree was traversed from tips to root, and the change along every branch was calculated by subtracting the ancestral state from the descendant state. Positive differences were counted toward gains, and negative differences were counted toward losses. Changes across all COGs were then summed separately for gains and losses and normalized by the total tree length to obtain gain and loss rates. The overall flux rate is the ratio of gain rate to loss rate.

### Data Availability

ATGC data that were used in this study is publicly available and can be found at https://ftp.ncbi.nih.gov/pub/wolf/ATGC/atgc-25/. Analyzed genome IDs and derived data, including *dN/dS* values, splittable ATGCs, decomposition results, simulated alignments, and decomposition script are deposited in Zenodo under https://doi.org/10.5281/zenodo.17575555.

## Notes

https://doi.org/10.5281/zenodo.17575555

### Competing Interest Statement

The authors have declared no competing interest.

### Funding

- National Institutes of Health: Intramural Research Program

