## Supplementary material for "The heterogeneous selection landscape of genome evolution in prokaryotes"

for

by

Roman Kogay[1], Svetlana Karamycheva[1], Nash D. Rochman[1,2,3] Yuri I. Wolf[1], Eugene V. Koonin[1]

[1]Computational Biology Branch, Division of Intramural Research, National Library of Medicine, National Institutes of Health, Bethesda, MD 20894, USA
[2] City University of New York Graduate School of Public Health and Health Policy, Department of Epidemiology and Biostatistics, New York, NY, United States
[3] Institute for Implementation Science in Population Health, City University of New York School of Public Health, New York, NY

### Supplementary Figures

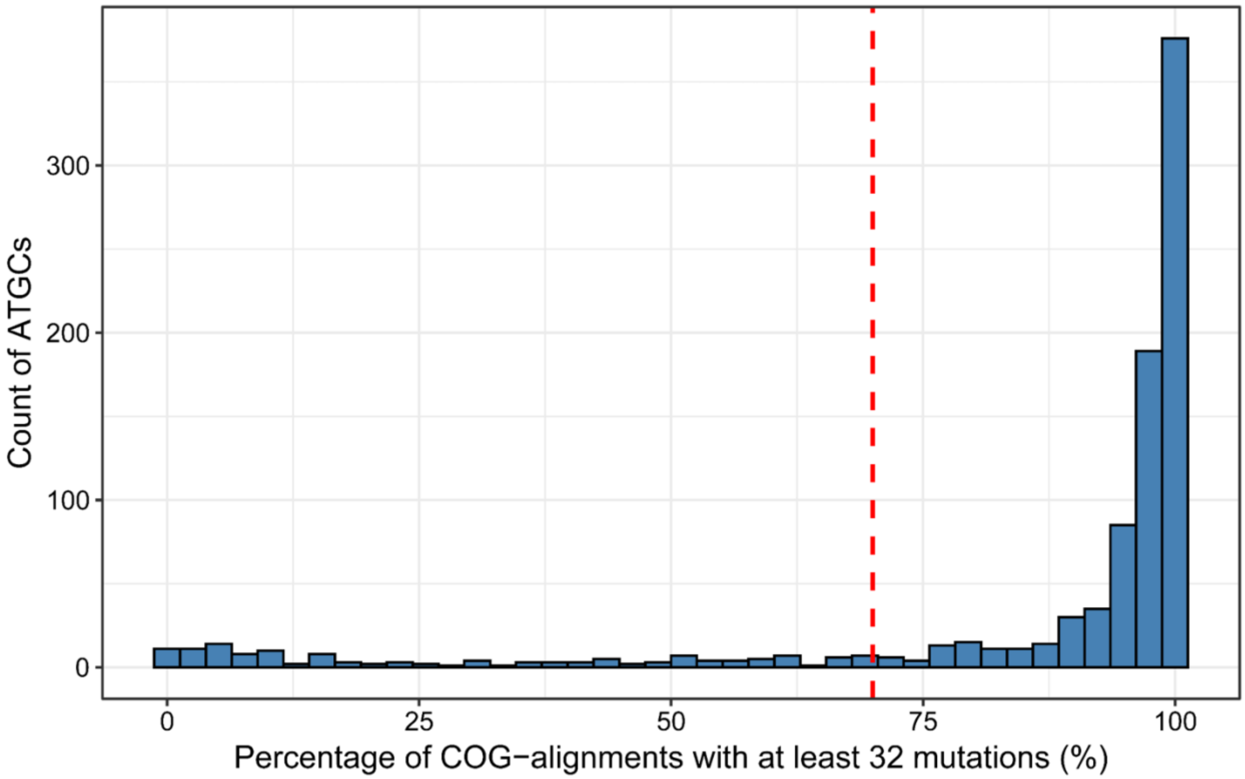


**Figure S1. Distribution of genetic diversity in alignments across ATGCs**. Histogram shows the distribution COG alignments with at least 32 mutations across ATGCs, whereas the dashed red line depicts the 70% cutoff that was used to filter out genetically homogeneous ATGCs.

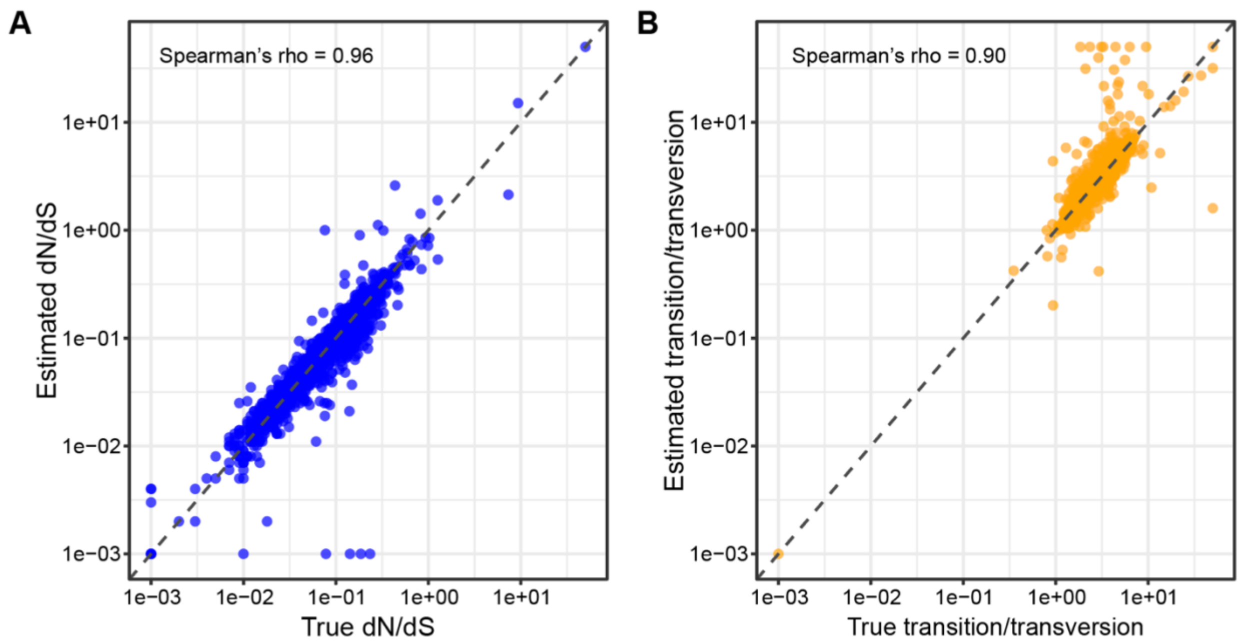


**Figure S2. Validation of (A) *dN/dS* and (B) transition/transversion estimation by IQTree across 2,724 simulated alignments**

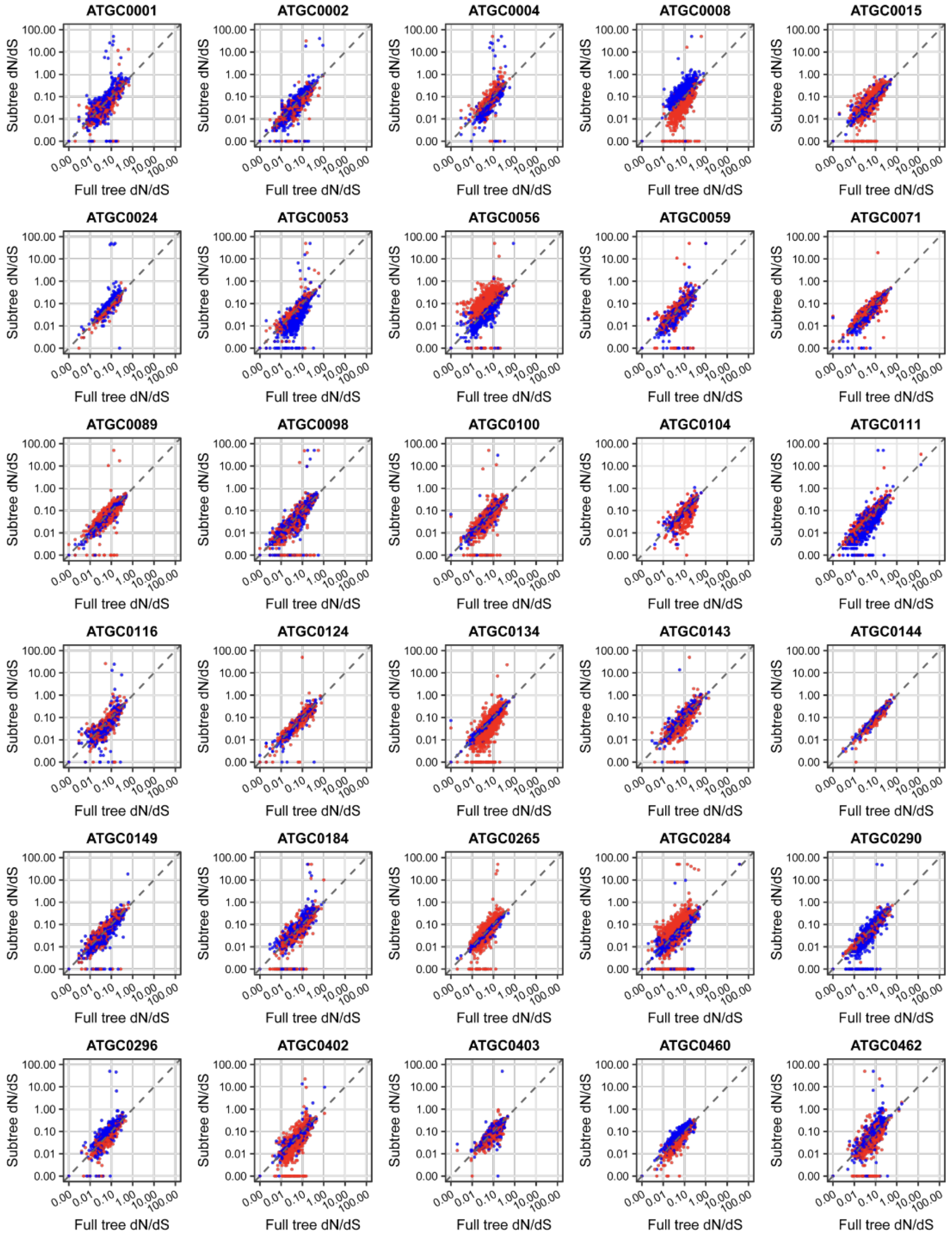


**Figure S3. Pairwise comparisons of *dN/dS* estimates between subtrees and full trees of 30 "splittable" ATGCs.** Each point represents *dN/dS* value of COG derived from two subtrees of the same ATGC (red and blue). The dashed line indicates a 1:1 relationship.

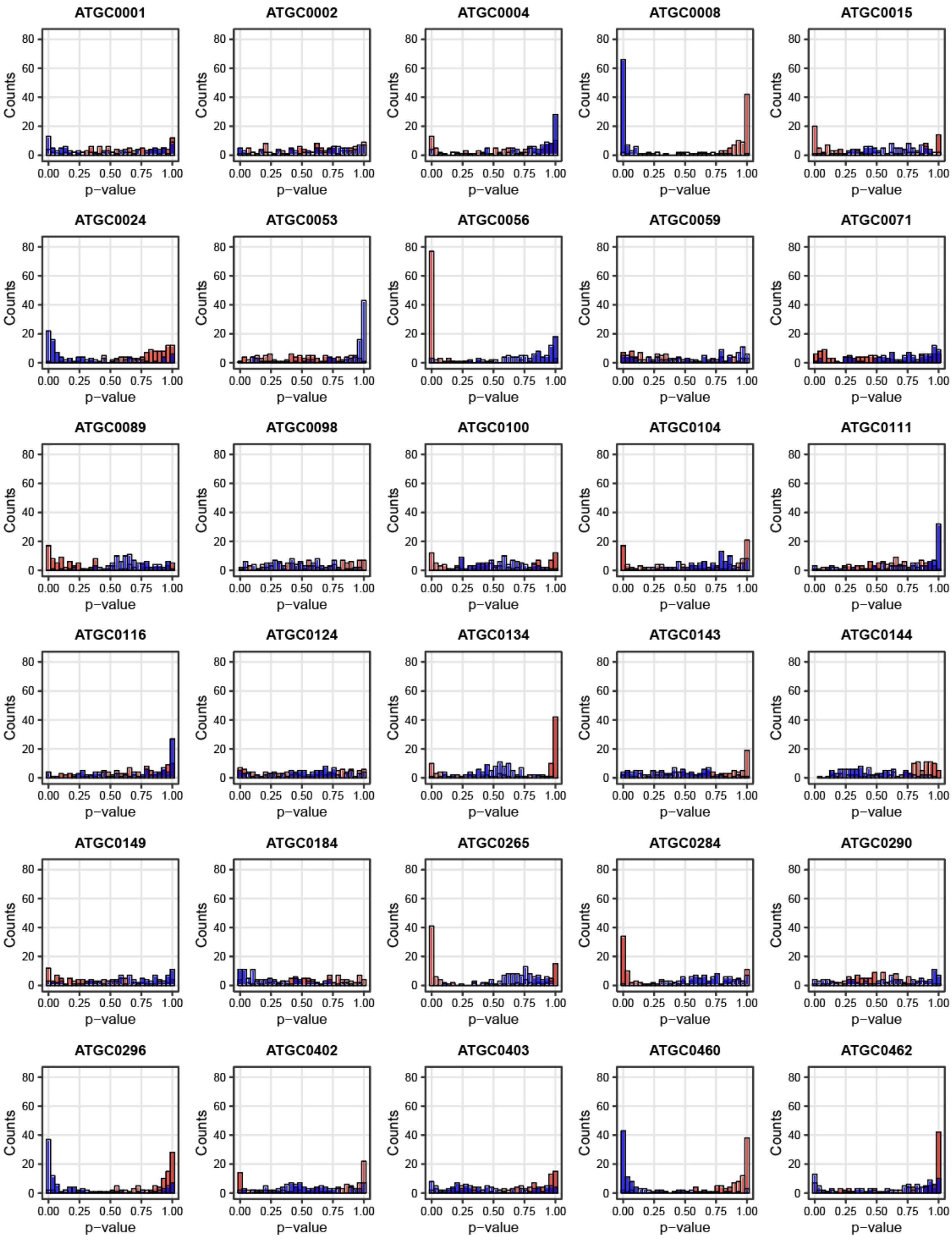


**Figure S4. Bootstrap analysis of interclade variation in splittable ATGCs.** Histograms show the distribution of p-values for two subtrees of each splittable ATGC (red and blue), based on

100 randomly selected COGs. For each subtree, the p-value represents the proportion of bootstrap replicates in which the subtree's *dN/dS* value exceeded that of the corresponding replicate derived from the full ATGC alignment. A p-value of 0 indicates that the subtree's *dN/dS* never exceeded the full-tree value, whereas a p-value of 1 indicates that it did so across all bootstrap replicates.

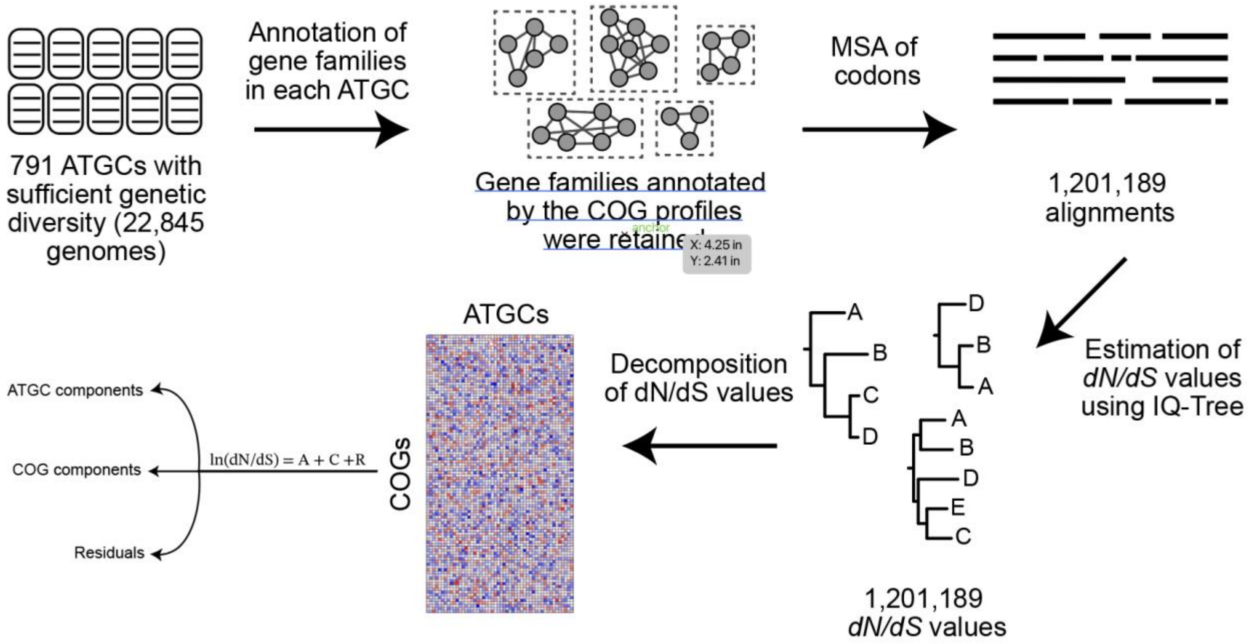


**Figure S5. Overview of the workflow for estimating lineage-specific and gene-specific selective constraints.**

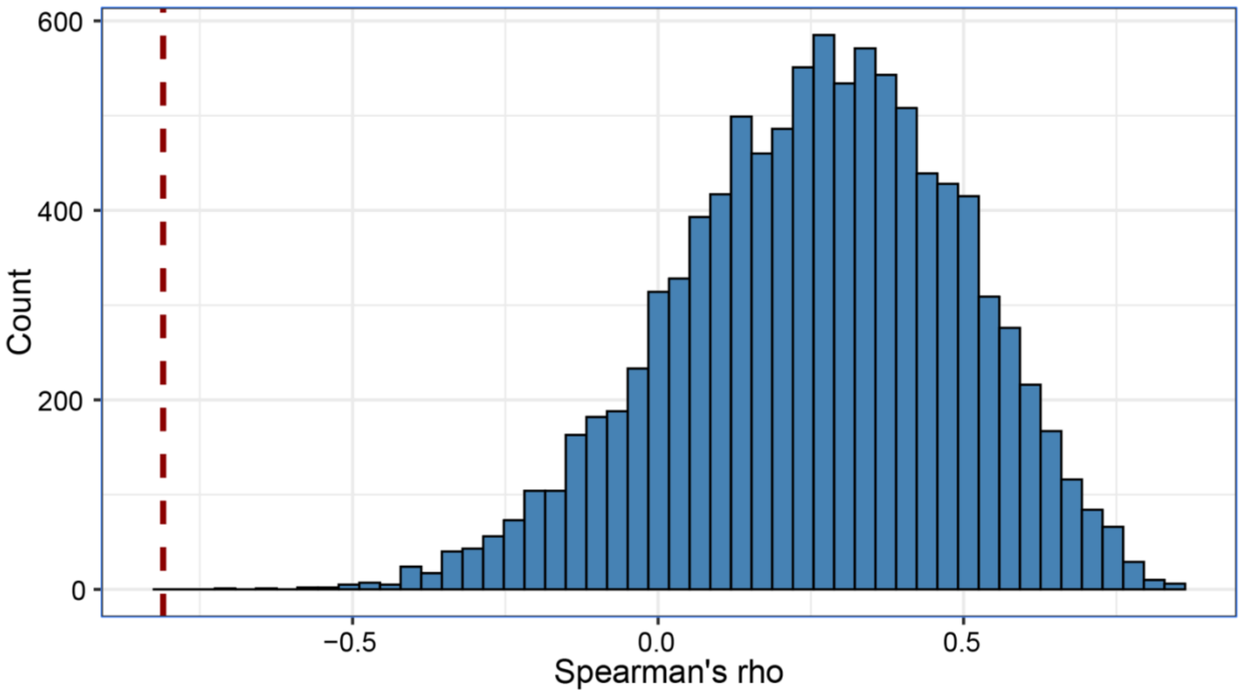


**Figure S6. Bootstrap test of correlation strength in Archaea versus Bacteria.** Histogram shows the distribution of Spearman's correlation coefficients between genome size and ATGC-specific constraint obtained from 10,000 random bootstrap resamples of 19 bacterial ATGCs. The dashed red line denotes the observed correlation for Archaea (Spearman's rho = -0.82).

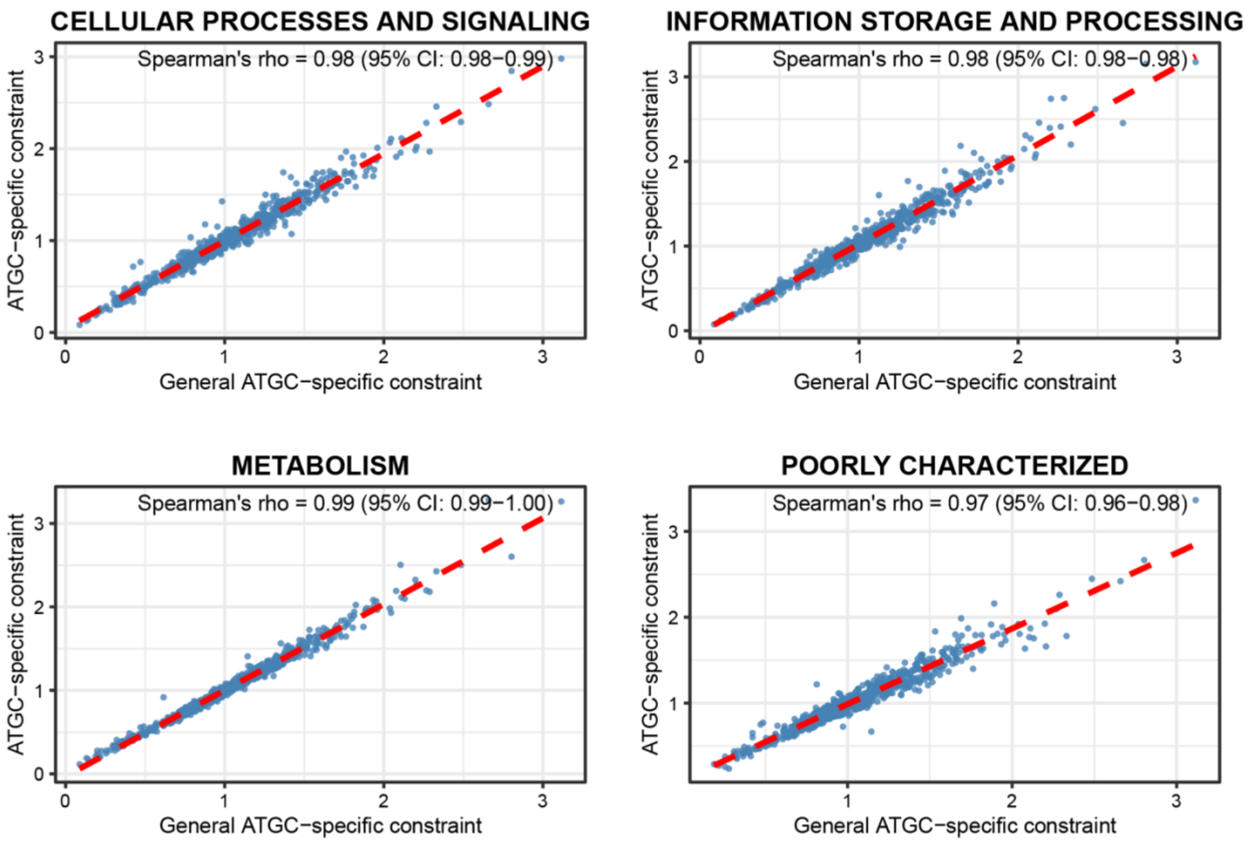


**Figure S7. Correlation between genome-wide and functional class ATGC-specific constraints.**

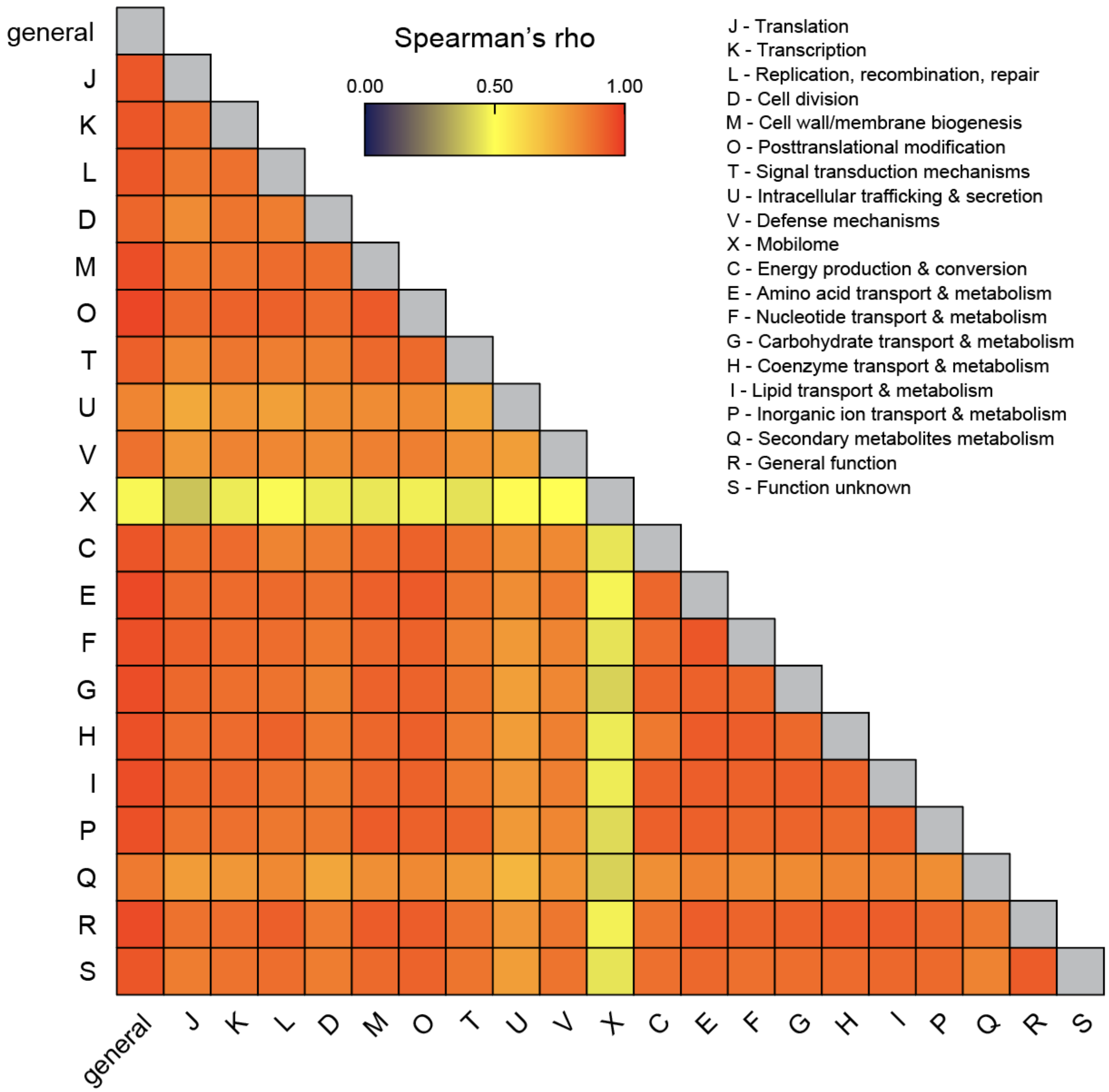


**Figure S8. Heatmap of correlations between ATGC-specific constraints across individual COG functional categories and genome-wide ('general') estimates.** Only ATGCs with *dN/dS* estimates available for at least 10 COGs were included in the analysis, and only COG functional groups represented by at least 500 ATGCs were retained for decomposition analyses. In all cases, the Spearman's correlation coefficients are statistically significant.

### Supplementary Tables

**Table S1. Summary of genome and evolutionary parameters estimated for 791 ATGCs.**

| ATGC ID | Genome Size | Constraint | Gain Rate | Loss Rate | Flux Rate |
|---|---|---|---|---|---|
| ATGC0001 | 4961446 | 1.00 | 32788.22 | 34692.33 | 0.95 |
| ATGC0002 | 5659878 | 1.15 | 40070.08 | 31550.28 | 1.27 |
| ATGC0003 | 2098294 | 0.85 | 5763.31 | 7697.84 | 0.75 |
| ATGC0004 | 1899037 | 0.67 | 4996.96 | 6686.70 | 0.75 |
| ATGC0005 | 2144862 | 0.84 | 5180.88 | 4659.26 | 1.11 |
| ATGC0007 | 2548542 | 0.78 | 12650.56 | 12291.49 | 1.03 |
| ATGC0008 | 1860070 | 0.70 | 16631.09 | 17388.03 | 0.96 |
| ATGC0009 | 1995667 | 0.90 | 6162.79 | 6491.00 | 0.95 |
| ATGC0010 | 2031692 | 0.45 | 9796.56 | 9084.50 | 1.08 |
| ATGC0012 | 2106259 | 1.40 | 7018.05 | 4423.48 | 1.59 |
| ATGC0014 | 5751681 | 0.78 | 35903.69 | 34402.46 | 1.04 |
| ATGC0015 | 4102385 | 0.79 | 17596.58 | 13545.55 | 1.30 |
| ATGC0016 | 3592666 | 0.69 | 9704.52 | 10357.78 | 0.94 |
| ATGC0017 | 3816932 | 0.69 | 14610.73 | 12065.18 | 1.21 |
| ATGC0018 | 4327255 | 0.67 | 64669.88 | 34003.03 | 1.90 |
| ATGC0019 | 5450197 | 0.77 | 61696.16 | 43951.76 | 1.40 |
| ATGC0020 | 3581994 | 0.49 | 26861.06 | 22596.60 | 1.19 |
| ATGC0021 | 1050220 | 0.75 | 1479.92 | 1518.51 | 0.97 |
| ATGC0022 | 1168962 | 0.42 | 2727.51 | 3828.40 | 0.71 |
| ATGC0024 | 4419298 | 0.67 | 11589.39 | 13628.92 | 0.85 |
| ATGC0028 | 6140828 | 0.84 | 7053.17 | 6437.50 | 1.10 |
| ATGC0030 | 5075529 | 1.18 | 18594.59 | 12907.01 | 1.44 |
| ATGC0031 | 6995345 | 1.20 | 5995.91 | 5679.72 | 1.06 |
| ATGC0032 | 977134 | 0.70 | 866.82 | 890.08 | 0.97 |
| ATGC0035 | 1049305 | 0.93 | 4717.61 | 4818.93 | 0.98 |
| ATGC0036 | 919490 | 0.81 | 288.89 | 282.60 | 1.02 |
| ATGC0037 | 985495 | 0.53 | 29464.64 | 43324.88 | 0.68 |
| ATGC0039 | 741809 | 0.98 | 1141.12 | 1054.73 | 1.08 |
| ATGC0044 | 1283764 | 0.25 | 21145.72 | 17690.84 | 1.20 |
| ATGC0046 | 1302786 | 0.22 | 18429.19 | 18806.17 | 0.98 |
| ATGC0047 | 1198154 | 0.57 | 949.80 | 1012.34 | 0.94 |
| ATGC0050 | 1625134 | 0.41 | 3445.76 | 3779.49 | 0.91 |
| ATGC0051 | 2196973 | 0.51 | 18166.67 | 15402.24 | 1.18 |
| ATGC0052 | 2819430 | 0.86 | 30349.14 | 26151.69 | 1.16 |
| ATGC0053 | 2552560 | 0.97 | 8393.99 | 7934.27 | 1.06 |

| | | | | | |
|---|---|---|---|---|---|
| ATGC0055 | 2605960 | 0.80 | 7881.75 | 5858.46 | 1.35 |
| ATGC0056 | 3039024 | 1.10 | 8322.10 | 8902.72 | 0.93 |
| ATGC0057 | 3310792 | 1.05 | 22915.35 | 21077.60 | 1.09 |
| ATGC0059 | 1966342 | 1.23 | 8554.03 | 7007.75 | 1.22 |
| ATGC0060 | 2102630 | 0.83 | 26344.59 | 18087.40 | 1.46 |
| ATGC0061 | 1910306 | 0.33 | 52844.87 | 57096.62 | 0.93 |
| ATGC0062 | 2098598 | 0.37 | 58714.67 | 50776.39 | 1.16 |
| ATGC0063 | 2111012 | 0.83 | 30979.18 | 22405.77 | 1.38 |
| ATGC0064 | 2551652 | 1.60 | 620.43 | 470.02 | 1.32 |
| ATGC0065 | 2594790 | 0.84 | 2781.36 | 2428.47 | 1.15 |
| ATGC0066 | 2145019 | 0.74 | 5148.27 | 5384.29 | 0.96 |
| ATGC0067 | 2370577 | 0.54 | 1625.09 | 1720.80 | 0.94 |
| ATGC0069 | 3282708 | 0.56 | 2266.27 | 2165.76 | 1.05 |
| ATGC0070 | 2368334 | 0.62 | 654.22 | 471.97 | 1.39 |
| ATGC0071 | 5994216 | 1.14 | 16432.57 | 12774.21 | 1.29 |
| ATGC0072 | 6704980 | 1.16 | 97441.76 | 41906.65 | 2.33 |
| ATGC0073 | 4563432 | 1.39 | 9993.36 | 6811.95 | 1.47 |
| ATGC0075 | 6179442 | 1.10 | 14929.18 | 13379.27 | 1.12 |
| ATGC0078 | 6479828 | 1.16 | 9837.56 | 11270.97 | 0.87 |
| ATGC0079 | 5238364 | 1.21 | 11653.93 | 10363.35 | 1.12 |
| ATGC0081 | 4089027 | 0.61 | 27762.36 | 21505.89 | 1.29 |
| ATGC0082 | 3347523 | 0.81 | 37177.35 | 29745.33 | 1.25 |
| ATGC0083 | 4642420 | 0.90 | 9744.48 | 7326.33 | 1.33 |
| ATGC0088 | 7154488 | 0.77 | 25342.19 | 31914.30 | 0.79 |
| ATGC0089 | 7526082 | 0.96 | 35178.25 | 47494.61 | 0.74 |
| ATGC0091 | 7241186 | 0.90 | 15344.67 | 10787.92 | 1.42 |
| ATGC0093 | 2655201 | 0.62 | 19273.66 | 21170.39 | 0.91 |
| ATGC0097 | 3834490 | 0.88 | 6439.87 | 5655.02 | 1.14 |
| ATGC0098 | 5232355 | 1.37 | 46580.69 | 30082.72 | 1.55 |
| ATGC0099 | 4908144 | 1.47 | 12401.61 | 11127.77 | 1.11 |
| ATGC0100 | 4962647 | 1.23 | 17893.12 | 16567.61 | 1.08 |
| ATGC0101 | 4714731 | 1.26 | 17118.16 | 14440.95 | 1.19 |
| ATGC0103 | 5472828 | 1.53 | 8481.69 | 6937.30 | 1.22 |
| ATGC0104 | 2326070 | 0.52 | 5591.38 | 6675.85 | 0.84 |
| ATGC0107 | 1674184 | 0.75 | 1719.92 | 1498.04 | 1.15 |
| ATGC0108 | 2954278 | 1.52 | 11722.46 | 7989.19 | 1.47 |
| ATGC0109 | 4088961 | 0.99 | 5105.49 | 5908.78 | 0.86 |
| ATGC0110 | 5389650 | 1.02 | 11092.85 | 8883.85 | 1.25 |
| ATGC0111 | 4808748 | 1.33 | 15016.58 | 11834.24 | 1.27 |
| ATGC0112 | 4388462 | 2.05 | 15878.49 | 11254.06 | 1.41 |

| | | | | | |
|---|---|---|---|---|---|
| ATGC0113 | 5073466 | 1.18 | 7149.30 | 4756.86 | 1.50 |
| ATGC0114 | 1887933 | 0.89 | 3375.77 | 2437.34 | 1.39 |
| ATGC0115 | 2340216 | 0.72 | 33656.07 | 20905.29 | 1.61 |
| ATGC0116 | 2524779 | 0.89 | 5101.44 | 5923.79 | 0.86 |
| ATGC0117 | 2145996 | 0.88 | 5449.83 | 4821.92 | 1.13 |
| ATGC0118 | 2326664 | 1.00 | 1024.52 | 1252.05 | 0.82 |
| ATGC0120 | 5002082 | 1.24 | 8237.53 | 5448.12 | 1.51 |
| ATGC0121 | 5278067 | 1.47 | 1366.40 | 760.90 | 1.80 |
| ATGC0122 | 6872475 | 0.81 | 16214.38 | 17123.27 | 0.95 |
| ATGC0123 | 6742135 | 0.92 | 15377.06 | 18009.45 | 0.85 |
| ATGC0124 | 6877215 | 0.85 | 16083.26 | 17970.33 | 0.89 |
| ATGC0126 | 5676673 | 1.00 | 16510.66 | 12135.92 | 1.36 |
| ATGC0130 | 3907473 | 0.88 | 9115.53 | 6521.69 | 1.40 |
| ATGC0133 | 3895750 | 0.76 | 7046.81 | 5142.31 | 1.37 |
| ATGC0134 | 5067146 | 1.00 | 19973.65 | 23251.13 | 0.86 |
| ATGC0135 | 4636005 | 1.22 | 12319.55 | 9273.40 | 1.33 |
| ATGC0136 | 3311836 | 0.93 | 13599.15 | 18939.77 | 0.72 |
| ATGC0137 | 2216603 | 0.65 | 6502.98 | 9034.01 | 0.72 |
| ATGC0138 | 1893705 | 0.74 | 3027.16 | 3771.67 | 0.80 |
| ATGC0139 | 7888977 | 0.83 | 18521.06 | 20239.53 | 0.92 |
| ATGC0141 | 8835682 | 0.76 | 19376.44 | 17933.99 | 1.08 |
| ATGC0143 | 1695084 | 0.64 | 7671.11 | 6317.33 | 1.21 |
| ATGC0144 | 905827 | 0.44 | 3728.37 | 2615.81 | 1.43 |
| ATGC0145 | 1328234 | 0.30 | 49940.25 | 17861.73 | 2.80 |
| ATGC0146 | 1860725 | 0.64 | 4014.58 | 4007.02 | 1.00 |
| ATGC0147 | 1746697 | 0.98 | 2727.64 | 1847.10 | 1.48 |
| ATGC0149 | 4019831 | 1.27 | 25617.70 | 23888.38 | 1.07 |
| ATGC0150 | 2093784 | 1.14 | 5219.72 | 3312.05 | 1.58 |
| ATGC0155 | 646449 | 0.37 | 244.60 | 173.24 | 1.41 |
| ATGC0157 | 3247995 | 1.24 | 18309.43 | 14590.58 | 1.25 |
| ATGC0159 | 2522885 | 0.70 | 5089.26 | 5924.97 | 0.86 |
| ATGC0160 | 2495938 | 0.91 | 5579.95 | 3170.95 | 1.76 |
| ATGC0161 | 2744312 | 1.48 | 3429.12 | 2635.49 | 1.30 |
| ATGC0162 | 2963612 | 0.67 | 100172.98 | 66980.48 | 1.50 |
| ATGC0163 | 3011798 | 0.55 | 113861.84 | 135556.88 | 0.84 |
| ATGC0165 | 4606604 | 1.01 | 7848.54 | 6444.29 | 1.22 |
| ATGC0166 | 4320012 | 0.93 | 4804.20 | 2577.86 | 1.86 |
| ATGC0167 | 4281685 | 0.93 | 5066.64 | 3666.12 | 1.38 |
| ATGC0168 | 2194207 | 0.84 | 16534.18 | 21677.09 | 0.76 |
| ATGC0169 | 2374850 | 0.73 | 9480.58 | 8548.94 | 1.11 |

| | | | | | |
|---|---|---|---|---|---|
| ATGC0173 | 5371816 | 0.91 | 6950.69 | 5344.66 | 1.30 |
| ATGC0174 | 3792700 | 0.94 | 8211.97 | 5188.69 | 1.58 |
| ATGC0177 | 1663938 | 0.73 | 1495.36 | 1027.02 | 1.46 |
| ATGC0180 | 1924674 | 0.61 | 2852.85 | 2254.22 | 1.27 |
| ATGC0181 | 2739610 | 0.68 | 8523.05 | 6169.91 | 1.38 |
| ATGC0182 | 2895001 | 0.69 | 18858.32 | 11617.80 | 1.62 |
| ATGC0183 | 4107424 | 1.06 | 66748.72 | 94868.90 | 0.70 |
| ATGC0184 | 3447086 | 0.71 | 19262.88 | 14875.40 | 1.29 |
| ATGC0185 | 4160292 | 0.53 | 12468.22 | 9755.16 | 1.28 |
| ATGC0187 | 2277260 | 1.44 | 20046.96 | 13338.42 | 1.50 |
| ATGC0188 | 5746272 | 0.92 | 60372.41 | 79647.56 | 0.76 |
| ATGC0189 | 6803198 | 0.99 | 20865.88 | 19279.37 | 1.08 |
| ATGC0190 | 4653851 | 1.45 | 6002.80 | 6931.69 | 0.87 |
| ATGC0192 | 1885386 | 0.71 | 2615.31 | 2394.79 | 1.09 |
| ATGC0193 | 2050591 | 0.77 | 4670.58 | 4734.68 | 0.99 |
| ATGC0194 | 3764261 | 0.92 | 14335.69 | 10211.69 | 1.40 |
| ATGC0196 | 2179788 | 1.05 | 15640.79 | 11172.92 | 1.40 |
| ATGC0199 | 5822107 | 0.83 | 23072.30 | 20944.61 | 1.10 |
| ATGC0201 | 1955442 | 0.46 | 2808.96 | 3430.88 | 0.82 |
| ATGC0204 | 6895682 | 0.73 | 12858.14 | 13301.20 | 0.97 |
| ATGC0206 | 4750213 | 0.79 | 29128.93 | 19991.78 | 1.46 |
| ATGC0207 | 3924697 | 0.67 | 15578.81 | 15475.69 | 1.01 |
| ATGC0210 | 1401460 | 0.68 | 7039.27 | 4553.74 | 1.55 |
| ATGC0211 | 5316784 | 0.95 | 9598.61 | 11739.79 | 0.82 |
| ATGC0212 | 3659940 | 0.91 | 10225.13 | 6453.15 | 1.58 |
| ATGC0213 | 5632330 | 0.97 | 14620.28 | 9902.26 | 1.48 |
| ATGC0216 | 5527842 | 1.11 | 12237.96 | 20650.92 | 0.59 |
| ATGC0219 | 2586443 | 1.32 | 27764.08 | 14096.76 | 1.97 |
| ATGC0220 | 3058643 | 1.22 | 1963.25 | 1470.74 | 1.33 |
| ATGC0222 | 4140668 | 1.11 | 8314.92 | 6863.70 | 1.21 |
| ATGC0223 | 638504 | 0.26 | 771.21 | 560.54 | 1.38 |
| ATGC0224 | 2637428 | 0.44 | 9370.81 | 10806.71 | 0.87 |
| ATGC0225 | 10246892 | 0.87 | 20001.63 | 17773.43 | 1.13 |
| ATGC0228 | 2178998 | 0.95 | 545.34 | 510.39 | 1.07 |
| ATGC0229 | 8529284 | 0.83 | 20616.85 | 27772.25 | 0.74 |
| ATGC0231 | 4360656 | 1.65 | 7966.19 | 5526.17 | 1.44 |
| ATGC0234 | 3294124 | 1.33 | 7238.77 | 6397.12 | 1.13 |
| ATGC0235 | 4351630 | 1.09 | 6766.42 | 6745.36 | 1.00 |
| ATGC0236 | 2837083 | 2.27 | 2971.19 | 1738.57 | 1.71 |
| ATGC0237 | 2379373 | 0.43 | 34095.82 | 28644.28 | 1.19 |

| | | | | | |
|---|---|---|---|---|---|
| ATGC0239 | 3877096 | 0.86 | 40936.89 | 32608.30 | 1.26 |
| ATGC0241 | 9289672 | 0.90 | 23342.42 | 20018.15 | 1.17 |
| ATGC0243 | 3208634 | 1.30 | 1797.51 | 1485.10 | 1.21 |
| ATGC0244 | 3194182 | 0.81 | 2565.10 | 2386.37 | 1.07 |
| ATGC0245 | 3071888 | 0.54 | 15457.87 | 10461.54 | 1.48 |
| ATGC0246 | 1230379 | 0.32 | 1572.62 | 1464.35 | 1.07 |
| ATGC0247 | 3782950 | 1.44 | 2421.79 | 1651.88 | 1.47 |
| ATGC0248 | 3762895 | 0.93 | 30583.37 | 26806.03 | 1.14 |
| ATGC0249 | 3408783 | 1.24 | 7535.07 | 5635.49 | 1.34 |
| ATGC0251 | 790674 | 0.32 | 230.47 | 233.34 | 0.99 |
| ATGC0257 | 3276390 | 0.84 | 4691.69 | 4238.99 | 1.11 |
| ATGC0261 | 2372880 | 0.92 | 10367.65 | 7905.50 | 1.31 |
| ATGC0263 | 1668548 | 1.35 | 532.06 | 325.38 | 1.64 |
| ATGC0265 | 1833375 | 1.10 | 2063.76 | 2033.78 | 1.01 |
| ATGC0270 | 2191533 | 2.29 | 3492.37 | 2243.63 | 1.56 |
| ATGC0272 | 4560446 | 0.50 | 14715.45 | 10435.48 | 1.41 |
| ATGC0277 | 3020235 | 0.80 | 5596.98 | 3249.38 | 1.72 |
| ATGC0280 | 2082088 | 1.64 | 7728.81 | 4241.23 | 1.82 |
| ATGC0284 | 4102900 | 1.39 | 18598.13 | 13296.21 | 1.40 |
| ATGC0285 | 4529188 | 1.82 | 7383.77 | 5757.94 | 1.28 |
| ATGC0286 | 4990764 | 1.37 | 17928.90 | 16919.48 | 1.06 |
| ATGC0289 | 5878550 | 1.50 | 39369.89 | 32720.36 | 1.20 |
| ATGC0290 | 3765752 | 0.97 | 14111.41 | 10801.74 | 1.31 |
| ATGC0291 | 4785706 | 0.98 | 11147.93 | 9961.31 | 1.12 |
| ATGC0293 | 2836999 | 1.08 | 6959.60 | 6017.61 | 1.16 |
| ATGC0294 | 3991226 | 0.87 | 3773.35 | 2606.50 | 1.45 |
| ATGC0296 | 2036353 | 0.80 | 4490.85 | 5177.20 | 0.87 |
| ATGC0300 | 6984680 | 1.25 | 17787.42 | 12945.32 | 1.37 |
| ATGC0301 | 4712992 | 1.10 | 3977.69 | 3689.42 | 1.08 |
| ATGC0304 | 2528077 | 0.93 | 19881.22 | 9143.04 | 2.17 |
| ATGC0305 | 2390967 | 1.05 | 2218.18 | 2113.50 | 1.05 |
| ATGC0307 | 2874062 | 1.21 | 11367.00 | 5921.55 | 1.92 |
| ATGC0308 | 2646794 | 1.94 | 4578.28 | 3654.44 | 1.25 |
| ATGC0309 | 1563032 | 1.04 | 3474.49 | 1624.14 | 2.14 |
| ATGC0310 | 1856156 | 0.80 | 16187.23 | 11874.73 | 1.36 |
| ATGC0312 | 1967004 | 1.02 | 4063.93 | 2957.57 | 1.37 |
| ATGC0313 | 7272014 | 0.85 | 41825.75 | 44102.86 | 0.95 |
| ATGC0314 | 9482704 | 1.07 | 30591.31 | 26656.59 | 1.15 |
| ATGC0315 | 8324904 | 0.89 | 19539.12 | 18171.71 | 1.08 |
| ATGC0317 | 8951748 | 1.40 | 21495.15 | 13308.48 | 1.62 |

| | | | | | |
|---|---|---|---|---|---|
| ATGC0318 | 5756799 | 0.86 | 8440.92 | 6344.23 | 1.33 |
| ATGC0320 | 6515693 | 0.94 | 11110.15 | 9496.08 | 1.17 |
| ATGC0321 | 2296570 | 1.73 | 6823.56 | 5274.23 | 1.29 |
| ATGC0322 | 1389513 | 1.38 | 2708.05 | 1773.19 | 1.53 |
| ATGC0324 | 712385 | 1.14 | 1935.11 | 992.10 | 1.95 |
| ATGC0325 | 807195 | 1.53 | 1140.98 | 658.95 | 1.73 |
| ATGC0328 | 897443 | 2.66 | 638.60 | 346.01 | 1.85 |
| ATGC0330 | 2521219 | 0.55 | 2074.22 | 2277.72 | 0.91 |
| ATGC0333 | 3173955 | 0.97 | 1329.60 | 973.36 | 1.37 |
| ATGC0336 | 4177197 | 1.22 | 7966.89 | 6591.27 | 1.21 |
| ATGC0337 | 4867548 | 1.06 | 3590.60 | 2935.92 | 1.22 |
| ATGC0338 | 5634694 | 1.50 | 2829.11 | 2418.77 | 1.17 |
| ATGC0339 | 3696846 | 0.91 | 5305.75 | 6013.56 | 0.88 |
| ATGC0340 | 6565735 | 1.18 | 13077.44 | 12013.14 | 1.09 |
| ATGC0342 | 2125626 | 0.79 | 1415.47 | 760.65 | 1.86 |
| ATGC0344 | 3807884 | 0.94 | 29783.07 | 16773.09 | 1.78 |
| ATGC0345 | 6148198 | 0.73 | 10719.50 | 8195.00 | 1.31 |
| ATGC0346 | 4579117 | 0.68 | 40064.27 | 25776.25 | 1.55 |
| ATGC0348 | 2863518 | 0.67 | 4621.12 | 3856.00 | 1.20 |
| ATGC0352 | 4685234 | 0.90 | 15776.48 | 19466.45 | 0.81 |
| ATGC0353 | 8274043 | 0.74 | 18160.24 | 21548.60 | 0.84 |
| ATGC0354 | 8472196 | 0.81 | 17189.66 | 18189.16 | 0.95 |
| ATGC0356 | 8223505 | 0.93 | 19010.00 | 15453.13 | 1.23 |
| ATGC0357 | 9547725 | 0.73 | 32217.28 | 25922.82 | 1.24 |
| ATGC0358 | 7070328 | 1.19 | 30985.43 | 31177.75 | 0.99 |
| ATGC0359 | 7105405 | 0.83 | 7631.68 | 7401.38 | 1.03 |
| ATGC0361 | 5621332 | 0.76 | 8073.29 | 5386.02 | 1.50 |
| ATGC0364 | 4709076 | 1.61 | 10628.94 | 6661.97 | 1.60 |
| ATGC0366 | 4745670 | 0.70 | 25143.88 | 14657.60 | 1.72 |
| ATGC0367 | 2909547 | 1.03 | 7966.82 | 9289.76 | 0.86 |
| ATGC0368 | 3618118 | 1.00 | 30353.47 | 20316.31 | 1.49 |
| ATGC0373 | 3377510 | 1.20 | 5896.54 | 4293.49 | 1.37 |
| ATGC0374 | 4679364 | 1.30 | 7348.25 | 6902.20 | 1.06 |
| ATGC0376 | 3727550 | 0.90 | 4435.34 | 3490.00 | 1.27 |
| ATGC0377 | 2715780 | 0.87 | 3146.24 | 1965.33 | 1.60 |
| ATGC0378 | 2957548 | 1.35 | 6576.85 | 4171.57 | 1.58 |
| ATGC0379 | 3638672 | 0.96 | 8022.08 | 5606.29 | 1.43 |
| ATGC0381 | 6798722 | 1.22 | 11177.05 | 7251.05 | 1.54 |
| ATGC0382 | 5685386 | 1.14 | 5087.44 | 3447.32 | 1.48 |
| ATGC0383 | 8841474 | 0.94 | 28238.05 | 30193.77 | 0.94 |

| | | | | | |
|---|---|---|---|---|---|
| ATGC0385 | 2115460 | 0.67 | 10812.51 | 6431.03 | 1.68 |
| ATGC0386 | 2034560 | 1.12 | 4745.84 | 2946.50 | 1.61 |
| ATGC0387 | 4577275 | 1.36 | 4331.26 | 2957.62 | 1.46 |
| ATGC0388 | 3824196 | 1.87 | 8482.98 | 6114.64 | 1.39 |
| ATGC0389 | 3305193 | 1.43 | 2678.13 | 1770.28 | 1.51 |
| ATGC0390 | 4984172 | 0.90 | 13550.33 | 8939.46 | 1.52 |
| ATGC0392 | 4373091 | 0.76 | 8054.79 | 6379.71 | 1.26 |
| ATGC0393 | 5585899 | 1.11 | 8111.61 | 6229.63 | 1.30 |
| ATGC0394 | 5513615 | 1.06 | 10794.26 | 9137.46 | 1.18 |
| ATGC0396 | 2399175 | 1.28 | 7039.24 | 4526.25 | 1.56 |
| ATGC0397 | 2779391 | 0.90 | 3891.06 | 2655.77 | 1.47 |
| ATGC0400 | 3261108 | 1.25 | 12279.39 | 10018.79 | 1.23 |
| ATGC0402 | 6433952 | 1.07 | 21005.54 | 21570.44 | 0.97 |
| ATGC0403 | 5458602 | 0.66 | 10491.12 | 9641.33 | 1.09 |
| ATGC0406 | 1907984 | 0.61 | 355.30 | 254.59 | 1.40 |
| ATGC0408 | 6561336 | 0.90 | 14975.37 | 12197.58 | 1.23 |
| ATGC0409 | 9472622 | 1.21 | 23270.63 | 21927.13 | 1.06 |
| ATGC0410 | 3638176 | 0.99 | 8430.25 | 4565.26 | 1.85 |
| ATGC0411 | 4181119 | 0.94 | 36953.30 | 27035.23 | 1.37 |
| ATGC0413 | 3168302 | 0.67 | 2957.34 | 3029.81 | 0.98 |
| ATGC0414 | 7110136 | 0.94 | 11769.23 | 9781.43 | 1.20 |
| ATGC0415 | 2415716 | 0.78 | 2974.00 | 1744.19 | 1.71 |
| ATGC0416 | 4379414 | 1.14 | 14603.48 | 9636.50 | 1.52 |
| ATGC0417 | 2434262 | 0.72 | 1598.29 | 1232.56 | 1.30 |
| ATGC0418 | 6680451 | 1.02 | 12666.42 | 9709.09 | 1.30 |
| ATGC0421 | 7014538 | 0.94 | 14996.63 | 12105.15 | 1.24 |
| ATGC0422 | 7390548 | 1.58 | 28894.09 | 21101.05 | 1.37 |
| ATGC0430 | 1780614 | 1.81 | 4518.32 | 3089.62 | 1.46 |
| ATGC0432 | 465712 | 0.43 | 571.66 | 383.00 | 1.49 |
| ATGC0451 | 4989538 | 1.32 | 19670.93 | 19055.64 | 1.03 |
| ATGC0452 | 5095054 | 1.10 | 35821.60 | 24048.23 | 1.49 |
| ATGC0453 | 4792334 | 0.99 | 17387.50 | 11829.10 | 1.47 |
| ATGC0454 | 3394565 | 0.83 | 4689.20 | 4070.74 | 1.15 |
| ATGC0455 | 1590458 | 0.90 | 1834.89 | 1297.17 | 1.41 |
| ATGC0456 | 2803335 | 0.69 | 8804.15 | 8414.14 | 1.05 |
| ATGC0457 | 4147522 | 1.04 | 5190.91 | 4486.69 | 1.16 |
| ATGC0459 | 4921682 | 1.13 | 5506.88 | 5243.46 | 1.05 |
| ATGC0460 | 5106030 | 1.03 | 5083.56 | 5551.22 | 0.92 |
| ATGC0461 | 1482279 | 0.21 | 17216.35 | 16731.86 | 1.03 |
| ATGC0462 | 2011032 | 0.88 | 3649.51 | 4341.16 | 0.84 |

| | | | | | |
|---|---|---|---|---|---|
| ATGC0463 | 4685386 | 1.28 | 9659.00 | 9840.60 | 0.98 |
| ATGC0464 | 4924702 | 0.91 | 20827.06 | 18628.25 | 1.12 |
| ATGC0465 | 4074222 | 1.40 | 28309.28 | 21438.66 | 1.32 |
| ATGC0466 | 4627770 | 1.47 | 11742.46 | 7254.50 | 1.62 |
| ATGC0467 | 3723098 | 1.24 | 7245.34 | 8072.28 | 0.90 |
| ATGC0468 | 4935337 | 1.19 | 17364.51 | 17765.74 | 0.98 |
| ATGC0469 | 4882630 | 1.40 | 12704.59 | 9986.37 | 1.27 |
| ATGC0470 | 5419421 | 1.53 | 6329.41 | 6173.22 | 1.03 |
| ATGC0471 | 7485129 | 0.82 | 20282.20 | 20861.03 | 0.97 |
| ATGC0472 | 4248142 | 1.00 | 6315.51 | 5264.34 | 1.20 |
| ATGC0473 | 3544508 | 1.32 | 6948.98 | 5658.48 | 1.23 |
| ATGC0474 | 2807142 | 0.57 | 4081.47 | 4464.23 | 0.91 |
| ATGC0475 | 1277820 | 0.87 | 1698.64 | 1508.33 | 1.13 |
| ATGC0476 | 3852411 | 1.45 | 5475.57 | 3647.56 | 1.50 |
| ATGC0477 | 4546882 | 0.94 | 5552.54 | 5535.58 | 1.00 |
| ATGC0478 | 2044097 | 1.25 | 7501.48 | 5504.24 | 1.36 |
| ATGC0479 | 4143808 | 1.52 | 10170.79 | 7558.02 | 1.35 |
| ATGC0480 | 2841718 | 0.75 | 8519.09 | 7954.01 | 1.07 |
| ATGC0481 | 2658717 | 0.72 | 2997.16 | 3019.54 | 0.99 |
| ATGC0482 | 2136444 | 0.71 | 3612.49 | 2186.78 | 1.65 |
| ATGC0483 | 3634864 | 1.42 | 5402.76 | 5218.15 | 1.04 |
| ATGC0484 | 1093116 | 0.31 | 6689.39 | 3225.24 | 2.07 |
| ATGC0485 | 1862338 | 0.77 | 6285.73 | 5930.23 | 1.06 |
| ATGC0486 | 2949392 | 0.79 | 8648.09 | 6109.26 | 1.42 |
| ATGC0487 | 4919668 | 1.12 | 8091.34 | 6851.60 | 1.18 |
| ATGC0488 | 9249796 | 0.88 | 31024.85 | 30564.31 | 1.02 |
| ATGC0489 | 5488077 | 0.85 | 11116.71 | 9179.57 | 1.21 |
| ATGC0490 | 4501378 | 1.68 | 12285.27 | 8455.48 | 1.45 |
| ATGC0491 | 4014743 | 0.74 | 4273.22 | 3264.35 | 1.31 |
| ATGC0492 | 2837190 | 1.15 | 32835.15 | 17110.64 | 1.92 |
| ATGC0493 | 5179960 | 0.71 | 63146.24 | 47252.91 | 1.34 |
| ATGC0494 | 4830299 | 1.20 | 70806.26 | 24816.11 | 2.85 |
| ATGC0496 | 190740 | 0.14 | 555.23 | 480.80 | 1.15 |
| ATGC0497 | 271421 | 0.09 | 651.45 | 1291.86 | 0.50 |
| ATGC0498 | 5582097 | 1.21 | 8166.52 | 6692.78 | 1.22 |
| ATGC0499 | 5539937 | 0.80 | 20953.37 | 17116.87 | 1.22 |
| ATGC0500 | 2716119 | 1.22 | 2319.05 | 2008.10 | 1.15 |
| ATGC0501 | 1932531 | 0.77 | 1253.56 | 1546.45 | 0.81 |
| ATGC0502 | 2501198 | 0.91 | 6016.20 | 5180.53 | 1.16 |
| ATGC0503 | 2314355 | 0.71 | 1629.96 | 1746.29 | 0.93 |

| | | | | | |
|---|---|---|---|---|---|
| ATGC0504 | 6123005 | 1.05 | 19380.46 | 16937.21 | 1.14 |
| ATGC0505 | 5461560 | 1.24 | 27666.39 | 24951.40 | 1.11 |
| ATGC0506 | 4565821 | 0.88 | 7974.73 | 7177.52 | 1.11 |
| ATGC0507 | 8828441 | 0.72 | 30038.96 | 26401.99 | 1.14 |
| ATGC0508 | 10125106 | 0.83 | 19756.94 | 17325.85 | 1.14 |
| ATGC0509 | 5770288 | 1.27 | 8891.29 | 7485.33 | 1.19 |
| ATGC0510 | 6194848 | 1.08 | 7956.85 | 7544.29 | 1.05 |
| ATGC0511 | 4642888 | 0.98 | 50652.32 | 39760.48 | 1.27 |
| ATGC0512 | 2414982 | 1.05 | 4014.03 | 2998.28 | 1.34 |
| ATGC0513 | 5303410 | 1.07 | 9043.54 | 7150.04 | 1.26 |
| ATGC0514 | 5259057 | 0.92 | 2731.94 | 2442.79 | 1.12 |
| ATGC0515 | 1943768 | 0.76 | 9455.17 | 6698.56 | 1.41 |
| ATGC0516 | 5056766 | 0.76 | 10790.58 | 6436.47 | 1.68 |
| ATGC0517 | 4647428 | 0.68 | 13246.69 | 14653.67 | 0.90 |
| ATGC0518 | 4187153 | 0.78 | 5244.38 | 4776.00 | 1.10 |
| ATGC0519 | 6143576 | 0.83 | 10986.56 | 9387.73 | 1.17 |
| ATGC0520 | 5362622 | 0.53 | 21259.97 | 16277.80 | 1.31 |
| ATGC0521 | 3281994 | 0.93 | 8418.10 | 5267.33 | 1.60 |
| ATGC0522 | 2416187 | 1.15 | 4858.42 | 3872.88 | 1.25 |
| ATGC0524 | 6223742 | 0.90 | 6929.27 | 6298.19 | 1.10 |
| ATGC0525 | 4576670 | 1.00 | 9230.17 | 7536.43 | 1.22 |
| ATGC0526 | 4651252 | 1.16 | 6848.74 | 5333.43 | 1.28 |
| ATGC0527 | 2301850 | 0.50 | 3667.71 | 2599.61 | 1.41 |
| ATGC0530 | 2275603 | 0.98 | 10804.44 | 7977.04 | 1.35 |
| ATGC0531 | 8110698 | 0.95 | 16872.10 | 15421.08 | 1.09 |
| ATGC0535 | 4556624 | 0.79 | 5256.62 | 5044.71 | 1.04 |
| ATGC0536 | 3307202 | 0.62 | 2713.07 | 2262.55 | 1.20 |
| ATGC0537 | 5386144 | 1.71 | 11766.38 | 8185.23 | 1.44 |
| ATGC0538 | 11328784 | 0.71 | 54313.31 | 59442.97 | 0.91 |
| ATGC0539 | 3764118 | 0.78 | 6940.88 | 6124.95 | 1.13 |
| ATGC0540 | 2754290 | 0.66 | 2466.72 | 1724.05 | 1.43 |
| ATGC0541 | 5103923 | 0.46 | 47630.05 | 36095.30 | 1.32 |
| ATGC0542 | 1111523 | 0.40 | 1815.21 | 3050.46 | 0.60 |
| ATGC0543 | 5427866 | 1.51 | 6724.73 | 5246.57 | 1.28 |
| ATGC0544 | 7078281 | 1.22 | 7731.33 | 8352.07 | 0.93 |
| ATGC0545 | 4196542 | 1.81 | 4941.36 | 3964.46 | 1.25 |
| ATGC0546 | 7111375 | 0.92 | 13582.46 | 12673.06 | 1.07 |
| ATGC0547 | 3734239 | 0.66 | 4711.49 | 4426.60 | 1.06 |
| ATGC0548 | 12022941 | 0.86 | 34771.01 | 32461.79 | 1.07 |
| ATGC0549 | 3282664 | 0.80 | 14661.35 | 11257.01 | 1.30 |

| | | | | | |
|---|---|---|---|---|---|
| ATGC0550 | 2944264 | 0.98 | 1955.46 | 1263.14 | 1.55 |
| ATGC0551 | 8262537 | 0.89 | 12156.29 | 11406.06 | 1.07 |
| ATGC0552 | 5416158 | 1.33 | 10503.82 | 8673.91 | 1.21 |
| ATGC0553 | 2536550 | 1.12 | 2694.73 | 2153.73 | 1.25 |
| ATGC0554 | 6393864 | 0.96 | 8939.24 | 6939.14 | 1.29 |
| ATGC0555 | 351863 | 0.22 | 239.84 | 209.74 | 1.14 |
| ATGC0556 | 8236870 | 0.81 | 17899.82 | 14763.65 | 1.21 |
| ATGC0557 | 6570934 | 0.92 | 14814.80 | 10417.48 | 1.42 |
| ATGC0558 | 4742384 | 2.20 | 8077.35 | 5158.59 | 1.57 |
| ATGC0559 | 4036106 | 1.02 | 3089.73 | 2877.13 | 1.07 |
| ATGC0560 | 7720132 | 1.03 | 7820.09 | 6648.93 | 1.18 |
| ATGC0563 | 4751181 | 1.42 | 13570.31 | 13070.24 | 1.04 |
| ATGC0564 | 5104519 | 0.62 | 39074.57 | 33852.34 | 1.15 |
| ATGC0565 | 3763593 | 1.49 | 6350.36 | 3447.04 | 1.84 |
| ATGC0566 | 853179 | 0.79 | 975.07 | 712.03 | 1.37 |
| ATGC0567 | 2717368 | 0.73 | 7000.24 | 5237.30 | 1.34 |
| ATGC0568 | 1734658 | 0.37 | 37995.42 | 37376.48 | 1.02 |
| ATGC0569 | 3628592 | 1.28 | 8790.08 | 9333.59 | 0.94 |
| ATGC0570 | 2132299 | 1.32 | 2025.28 | 1520.26 | 1.33 |
| ATGC0571 | 8653730 | 0.51 | 40276.37 | 27266.32 | 1.48 |
| ATGC0572 | 7716609 | 1.22 | 11651.88 | 11380.39 | 1.02 |
| ATGC0573 | 4790868 | 1.24 | 5863.44 | 6559.43 | 0.89 |
| ATGC0574 | 5930964 | 0.68 | 10551.22 | 7906.46 | 1.33 |
| ATGC0575 | 5005459 | 1.62 | 8027.66 | 5806.09 | 1.38 |
| ATGC0576 | 465742 | 0.35 | 117.45 | 128.32 | 0.92 |
| ATGC0577 | 613221 | 0.32 | 782.98 | 599.54 | 1.31 |
| ATGC0578 | 4807151 | 0.96 | 7952.93 | 6311.29 | 1.26 |
| ATGC0580 | 6717976 | 1.55 | 27279.69 | 20052.39 | 1.36 |
| ATGC0581 | 3971838 | 0.81 | 5560.47 | 4550.67 | 1.22 |
| ATGC0582 | 641328 | 0.41 | 178.34 | 116.07 | 1.54 |
| ATGC0583 | 2459210 | 0.81 | 5801.44 | 4293.55 | 1.35 |
| ATGC0584 | 1842238 | 0.65 | 1360.93 | 1017.03 | 1.34 |
| ATGC0585 | 6352364 | 1.42 | 11826.93 | 8296.42 | 1.43 |
| ATGC0586 | 3563508 | 0.74 | 3008.89 | 2168.65 | 1.39 |
| ATGC0587 | 3617198 | 0.92 | 3582.72 | 2888.57 | 1.24 |
| ATGC0588 | 2336942 | 0.89 | 5067.60 | 3867.04 | 1.31 |
| ATGC0589 | 7673653 | 0.88 | 12318.35 | 10492.38 | 1.17 |
| ATGC0590 | 2529018 | 0.46 | 966.53 | 576.82 | 1.68 |
| ATGC0591 | 2280350 | 0.77 | 2903.53 | 1997.25 | 1.45 |
| ATGC0592 | 4273636 | 0.50 | 6407.93 | 4209.47 | 1.52 |

| | | | | | |
|---|---|---|---|---|---|
| ATGC0595 | 1081404 | 0.75 | 2036.51 | 1093.02 | 1.86 |
| ATGC0596 | 2725214 | 0.68 | 15745.09 | 12354.74 | 1.27 |
| ATGC0597 | 3104561 | 0.89 | 5671.97 | 4118.94 | 1.38 |
| ATGC0599 | 1377933 | 0.40 | 32744.60 | 25157.65 | 1.30 |
| ATGC0600 | 2272942 | 0.81 | 1991.66 | 1784.51 | 1.12 |
| ATGC0601 | 1561431 | 0.39 | 2761.22 | 1929.49 | 1.43 |
| ATGC0602 | 5335236 | 0.94 | 12658.01 | 11528.90 | 1.10 |
| ATGC0603 | 3504888 | 0.89 | 3210.94 | 2912.65 | 1.10 |
| ATGC0604 | 4926765 | 0.76 | 5087.59 | 3867.81 | 1.32 |
| ATGC0605 | 4704737 | 0.85 | 6582.79 | 6495.13 | 1.01 |
| ATGC0606 | 5349645 | 0.73 | 5604.11 | 4733.79 | 1.18 |
| ATGC0607 | 4208346 | 0.86 | 4816.72 | 4095.34 | 1.18 |
| ATGC0608 | 4786552 | 0.94 | 4215.36 | 3042.44 | 1.39 |
| ATGC0609 | 3340250 | 0.81 | 6253.40 | 4094.52 | 1.53 |
| ATGC0611 | 1676668 | 0.95 | 8105.70 | 6677.27 | 1.21 |
| ATGC0612 | 2060749 | 0.92 | 15566.57 | 11488.64 | 1.35 |
| ATGC0613 | 4164094 | 0.84 | 9477.70 | 9291.55 | 1.02 |
| ATGC0614 | 4548320 | 2.08 | 14005.24 | 11032.22 | 1.27 |
| ATGC0615 | 3636562 | 1.51 | 3058.84 | 3090.41 | 0.99 |
| ATGC0616 | 1294232 | 0.33 | 15496.29 | 15378.28 | 1.01 |
| ATGC0617 | 4527698 | 1.91 | 4670.28 | 5175.06 | 0.90 |
| ATGC0618 | 5834304 | 1.40 | 3485.70 | 2610.53 | 1.34 |
| ATGC0619 | 5078349 | 1.34 | 14646.46 | 9110.45 | 1.61 |
| ATGC0620 | 3973645 | 0.62 | 35781.35 | 26400.99 | 1.36 |
| ATGC0621 | 3121500 | 0.77 | 3399.13 | 2294.05 | 1.48 |
| ATGC0622 | 2747629 | 0.89 | 3407.82 | 2393.20 | 1.42 |
| ATGC0623 | 2760158 | 1.47 | 4808.44 | 3727.89 | 1.29 |
| ATGC0624 | 4782835 | 1.04 | 8871.05 | 7099.13 | 1.25 |
| ATGC0625 | 3301706 | 0.87 | 7987.74 | 5044.07 | 1.58 |
| ATGC0626 | 2500831 | 0.61 | 975.47 | 677.50 | 1.44 |
| ATGC0627 | 2734414 | 1.09 | 2421.24 | 1706.99 | 1.42 |
| ATGC0628 | 4520065 | 1.41 | 1923.32 | 1354.46 | 1.42 |
| ATGC0629 | 1889132 | 0.74 | 1478.60 | 990.13 | 1.49 |
| ATGC0630 | 445540 | 0.34 | 48.51 | 60.67 | 0.80 |
| ATGC0631 | 6862428 | 0.90 | 6328.61 | 5994.21 | 1.06 |
| ATGC0636 | 2127051 | 0.36 | 31267.74 | 22997.83 | 1.36 |
| ATGC0637 | 1850487 | 1.36 | 2300.52 | 1372.89 | 1.68 |
| ATGC0639 | 5946826 | 1.13 | 18913.54 | 13262.59 | 1.43 |
| ATGC0641 | 4195186 | 1.29 | 5723.44 | 8185.32 | 0.70 |
| ATGC0642 | 1660278 | 0.77 | 3520.67 | 2560.04 | 1.38 |

| | | | | | |
|---|---|---|---|---|---|
| ATGC0643 | 8270461 | 0.93 | 20634.00 | 16539.50 | 1.25 |
| ATGC0644 | 3825112 | 0.86 | 7553.68 | 4812.87 | 1.57 |
| ATGC0645 | 8068143 | 1.06 | 15074.59 | 11369.39 | 1.33 |
| ATGC0646 | 4338868 | 1.11 | 7060.54 | 4610.08 | 1.53 |
| ATGC0647 | 2744227 | 1.07 | 1444.66 | 1137.79 | 1.27 |
| ATGC0648 | 4669365 | 1.44 | 5188.67 | 5247.65 | 0.99 |
| ATGC0649 | 6922031 | 1.00 | 9007.95 | 7031.47 | 1.28 |
| ATGC0650 | 2243615 | 0.64 | 2928.95 | 2009.87 | 1.46 |
| ATGC0651 | 3380543 | 1.05 | 8800.40 | 6234.34 | 1.41 |
| ATGC0652 | 1344614 | 1.18 | 743.65 | 546.89 | 1.36 |
| ATGC0653 | 4090129 | 0.96 | 6006.92 | 3427.93 | 1.75 |
| ATGC0654 | 4204657 | 0.83 | 10101.90 | 7076.19 | 1.43 |
| ATGC0655 | 6641645 | 0.77 | 11059.04 | 7605.18 | 1.45 |
| ATGC0656 | 4370890 | 1.31 | 2653.38 | 2193.43 | 1.21 |
| ATGC0657 | 3459201 | 0.85 | 5034.19 | 4938.01 | 1.02 |
| ATGC0658 | 4674473 | 0.67 | 10900.36 | 6095.36 | 1.79 |
| ATGC0662 | 6112670 | 1.58 | 33357.63 | 23229.09 | 1.44 |
| ATGC0664 | 3361142 | 1.22 | 8250.89 | 5776.52 | 1.43 |
| ATGC0665 | 3736407 | 1.31 | 40328.22 | 17795.35 | 2.27 |
| ATGC0668 | 4873239 | 2.11 | 14786.33 | 9318.33 | 1.59 |
| ATGC0669 | 5242548 | 1.58 | 8114.25 | 6442.45 | 1.26 |
| ATGC0670 | 6907772 | 1.27 | 25265.45 | 19607.03 | 1.29 |
| ATGC0671 | 2409765 | 0.84 | 8273.59 | 5217.45 | 1.59 |
| ATGC0672 | 3404760 | 1.00 | 4125.58 | 2737.27 | 1.51 |
| ATGC0673 | 5344972 | 0.84 | 12854.80 | 8363.21 | 1.54 |
| ATGC0674 | 4464988 | 1.19 | 10661.86 | 7946.22 | 1.34 |
| ATGC0676 | 2360808 | 1.43 | 2681.45 | 1908.58 | 1.40 |
| ATGC0677 | 2011862 | 1.02 | 1594.89 | 1235.95 | 1.29 |
| ATGC0678 | 7166266 | 0.98 | 22060.52 | 13269.88 | 1.66 |
| ATGC0679 | 2876572 | 1.15 | 5695.21 | 4095.41 | 1.39 |
| ATGC0680 | 4348337 | 1.42 | 2188.67 | 1355.60 | 1.61 |
| ATGC0681 | 4655991 | 0.83 | 16077.95 | 12441.89 | 1.29 |
| ATGC0682 | 2853072 | 0.93 | 6539.27 | 3835.73 | 1.70 |
| ATGC0683 | 6633666 | 0.91 | 15313.71 | 12722.85 | 1.20 |
| ATGC0684 | 5117558 | 1.66 | 7398.76 | 7310.89 | 1.01 |
| ATGC0685 | 4507116 | 1.14 | 6042.18 | 4302.26 | 1.40 |
| ATGC0686 | 6135478 | 1.09 | 9353.93 | 8981.11 | 1.04 |
| ATGC0687 | 9059812 | 0.89 | 22675.43 | 17484.80 | 1.30 |
| ATGC0688 | 169219 | 0.47 | 102.58 | 86.98 | 1.18 |
| ATGC0689 | 2374864 | 0.86 | 1777.01 | 1605.71 | 1.11 |

| | | | | | |
|---|---|---|---|---|---|
| ATGC0690 | 4395968 | 0.77 | 3350.10 | 2351.30 | 1.42 |
| ATGC0691 | 3893206 | 0.69 | 2765.48 | 2385.68 | 1.16 |
| ATGC0692 | 3951517 | 1.18 | 6190.09 | 4232.65 | 1.46 |
| ATGC0693 | 3180673 | 1.64 | 4351.68 | 3233.96 | 1.35 |
| ATGC0694 | 3450279 | 1.42 | 4496.51 | 2763.96 | 1.63 |
| ATGC0699 | 3827531 | 1.96 | 27326.48 | 5071.53 | 5.39 |
| ATGC0700 | 2599652 | 1.05 | 19017.02 | 12609.01 | 1.51 |
| ATGC0702 | 4851806 | 1.21 | 18577.37 | 14135.04 | 1.31 |
| ATGC0703 | 2314753 | 1.36 | 2342.13 | 1610.00 | 1.45 |
| ATGC0704 | 4041318 | 1.27 | 11428.46 | 10914.07 | 1.05 |
| ATGC0705 | 4141447 | 2.48 | 9080.85 | 5603.97 | 1.62 |
| ATGC0706 | 6224392 | 1.43 | 16757.31 | 11228.00 | 1.49 |
| ATGC0707 | 6612542 | 0.69 | 27436.39 | 18848.17 | 1.46 |
| ATGC0708 | 5230650 | 1.56 | 17679.79 | 13491.79 | 1.31 |
| ATGC0709 | 4018990 | 1.11 | 14929.02 | 12767.42 | 1.17 |
| ATGC0710 | 4996294 | 0.97 | 22916.53 | 14108.31 | 1.62 |
| ATGC0711 | 8027492 | 1.21 | 10951.74 | 7792.23 | 1.41 |
| ATGC0712 | 6607828 | 1.12 | 16534.05 | 11672.42 | 1.42 |
| ATGC0713 | 6661285 | 1.73 | 20605.18 | 14387.72 | 1.43 |
| ATGC0714 | 723970 | 0.19 | 16075.38 | 7970.99 | 2.02 |
| ATGC0715 | 3362333 | 1.14 | 2429.14 | 1796.16 | 1.35 |
| ATGC0716 | 3584812 | 0.98 | 5109.84 | 3879.58 | 1.32 |
| ATGC0717 | 2182100 | 1.10 | 2186.59 | 1465.27 | 1.49 |
| ATGC0718 | 919993 | 2.11 | 1133.01 | 660.79 | 1.71 |
| ATGC0719 | 3424893 | 1.26 | 2470.53 | 1724.57 | 1.43 |
| ATGC0720 | 4839879 | 1.17 | 4469.00 | 3671.23 | 1.22 |
| ATGC0722 | 3827697 | 1.15 | 3941.87 | 3096.11 | 1.27 |
| ATGC0723 | 4255835 | 1.70 | 6363.35 | 5263.39 | 1.21 |
| ATGC0724 | 6147290 | 1.06 | 4373.11 | 3057.29 | 1.43 |
| ATGC0725 | 9240166 | 0.86 | 21594.01 | 17108.74 | 1.26 |
| ATGC0726 | 1986154 | 0.96 | 539.80 | 376.93 | 1.43 |
| ATGC0727 | 7424241 | 0.82 | 21798.11 | 16273.71 | 1.34 |
| ATGC0728 | 1796606 | 1.49 | 1946.67 | 1210.60 | 1.61 |
| ATGC0729 | 3979593 | 0.90 | 3475.90 | 2690.77 | 1.29 |
| ATGC0730 | 4876443 | 1.95 | 5992.24 | 3723.96 | 1.61 |
| ATGC0731 | 3370341 | 1.75 | 2521.58 | 1855.89 | 1.36 |
| ATGC0732 | 2496444 | 0.63 | 3498.27 | 2296.04 | 1.52 |
| ATGC0733 | 6049186 | 0.82 | 8886.74 | 8833.99 | 1.01 |
| ATGC0734 | 4376059 | 1.25 | 6728.30 | 4189.15 | 1.61 |
| ATGC0735 | 2702994 | 0.73 | 7145.83 | 4696.07 | 1.52 |

| | | | | | |
|---|---|---|---|---|---|
| ATGC0736 | 2287213 | 1.16 | 1656.62 | 932.06 | 1.78 |
| ATGC0737 | 5697957 | 1.46 | 10760.10 | 7851.23 | 1.37 |
| ATGC0738 | 3684270 | 1.45 | 5743.18 | 4978.53 | 1.15 |
| ATGC0739 | 4378193 | 1.58 | 6019.08 | 4071.18 | 1.48 |
| ATGC0740 | 4491948 | 0.91 | 4497.66 | 3552.20 | 1.27 |
| ATGC0745 | 6415420 | 1.76 | 26851.97 | 14588.57 | 1.84 |
| ATGC0746 | 4214921 | 1.38 | 14304.01 | 10388.98 | 1.38 |
| ATGC0750 | 6101332 | 2.80 | 34355.21 | 28354.98 | 1.21 |
| ATGC0752 | 3312578 | 0.52 | 21796.34 | 13517.78 | 1.61 |
| ATGC0755 | 2897281 | 1.21 | 2708.37 | 1656.65 | 1.63 |
| ATGC0757 | 2376036 | 1.14 | 3376.28 | 2448.31 | 1.38 |
| ATGC0758 | 2633957 | 0.82 | 6180.34 | 3292.52 | 1.88 |
| ATGC0759 | 4083035 | 1.29 | 7297.43 | 6571.68 | 1.11 |
| ATGC0760 | 2509910 | 1.10 | 3049.74 | 2069.54 | 1.47 |
| ATGC0762 | 1459038 | 1.11 | 919.47 | 653.68 | 1.41 |
| ATGC0763 | 3355004 | 1.52 | 8141.60 | 5088.35 | 1.60 |
| ATGC0764 | 635111 | 0.40 | 1556.81 | 1120.01 | 1.39 |
| ATGC0766 | 1753385 | 0.56 | 11923.31 | 7906.38 | 1.51 |
| ATGC0767 | 4657691 | 1.34 | 10882.00 | 7820.23 | 1.39 |
| ATGC0768 | 713367 | 0.95 | 5829.20 | 3861.57 | 1.51 |
| ATGC0769 | 3542126 | 1.80 | 3669.55 | 2510.72 | 1.46 |
| ATGC0770 | 3481664 | 1.53 | 11168.81 | 8071.60 | 1.38 |
| ATGC0771 | 5226988 | 1.21 | 12464.01 | 8782.34 | 1.42 |
| ATGC0772 | 140963 | 0.20 | 148.70 | 184.31 | 0.81 |
| ATGC0773 | 2287249 | 1.06 | 3725.29 | 2218.44 | 1.68 |
| ATGC0774 | 779363 | 0.37 | 159.45 | 233.90 | 0.68 |
| ATGC0775 | 4918994 | 1.56 | 7496.06 | 4832.85 | 1.55 |
| ATGC0776 | 8881052 | 1.03 | 17761.32 | 11698.53 | 1.52 |
| ATGC0777 | 3923376 | 1.04 | 10152.57 | 6498.69 | 1.56 |
| ATGC0778 | 5688986 | 0.73 | 29385.80 | 19666.87 | 1.49 |
| ATGC0779 | 5047056 | 1.79 | 4865.07 | 2975.58 | 1.64 |
| ATGC0780 | 3864079 | 0.92 | 4937.95 | 3599.31 | 1.37 |
| ATGC0781 | 5826844 | 1.15 | 14683.95 | 10701.14 | 1.37 |
| ATGC0782 | 2784822 | 0.94 | 1812.24 | 1179.31 | 1.54 |
| ATGC0783 | 5014524 | 1.15 | 6817.66 | 4988.55 | 1.37 |
| ATGC0784 | 2980819 | 1.05 | 3831.33 | 2516.81 | 1.52 |
| ATGC0785 | 1509068 | 1.10 | 1980.17 | 1208.51 | 1.64 |
| ATGC0786 | 152118 | 0.13 | 270.63 | 211.68 | 1.28 |
| ATGC0787 | 3954950 | 1.05 | 6742.14 | 4499.61 | 1.50 |
| ATGC0789 | 4915216 | 1.15 | 4570.24 | 2993.51 | 1.53 |

| | | | | | |
|---|---|---|---|---|---|
| ATGC0790 | 1603582 | 0.87 | 1890.06 | 1047.05 | 1.81 |
| ATGC0791 | 8577674 | 1.09 | 3329.51 | 2385.30 | 1.40 |
| ATGC0792 | 6669322 | 0.87 | 15853.19 | 10936.78 | 1.45 |
| ATGC0793 | 5089458 | 1.07 | 5242.05 | 3790.96 | 1.38 |
| ATGC0794 | 2548057 | 0.93 | 2945.29 | 1577.08 | 1.87 |
| ATGC0795 | 4120638 | 2.33 | 4001.60 | 2932.64 | 1.36 |
| ATGC0796 | 3844970 | 0.64 | 15516.57 | 9915.86 | 1.56 |
| ATGC0797 | 6038404 | 1.04 | 6840.44 | 3968.46 | 1.72 |
| ATGC0798 | 235831 | 0.32 | 84.23 | 79.45 | 1.06 |
| ATGC0799 | 4519234 | 2.04 | 7171.09 | 3890.64 | 1.84 |
| ATGC0800 | 7100882 | 0.73 | 18972.52 | 13053.88 | 1.45 |
| ATGC0801 | 8055216 | 1.01 | 12591.25 | 9745.04 | 1.29 |
| ATGC0802 | 3967624 | 1.15 | 7502.62 | 6246.85 | 1.20 |
| ATGC0803 | 5705613 | 1.21 | 7038.48 | 5280.38 | 1.33 |
| ATGC0805 | 3965954 | 0.99 | 3499.75 | 1763.76 | 1.98 |
| ATGC0806 | 5896700 | 0.78 | 7463.01 | 5444.66 | 1.37 |
| ATGC0807 | 6732154 | 0.69 | 20288.55 | 13110.74 | 1.55 |
| ATGC0808 | 651313 | 0.47 | 161.22 | 135.97 | 1.19 |
| ATGC0809 | 4026314 | 0.83 | 12267.57 | 9685.41 | 1.27 |
| ATGC0810 | 6263730 | 0.83 | 7289.21 | 5392.78 | 1.35 |
| ATGC0811 | 3269948 | 0.49 | 14260.27 | 9827.61 | 1.45 |
| ATGC0812 | 4503894 | 1.17 | 3800.83 | 2723.88 | 1.40 |
| ATGC0813 | 5984850 | 1.37 | 8957.93 | 6883.27 | 1.30 |
| ATGC0814 | 9285790 | 0.89 | 23989.33 | 17075.27 | 1.40 |
| ATGC0815 | 1456516 | 0.88 | 11040.93 | 8997.50 | 1.23 |
| ATGC0816 | 7884676 | 0.93 | 15699.33 | 9858.83 | 1.59 |
| ATGC0817 | 4738752 | 0.89 | 4766.90 | 3098.33 | 1.54 |
| ATGC0818 | 1524413 | 0.78 | 1785.30 | 1212.51 | 1.47 |
| ATGC0819 | 2714388 | 0.76 | 3410.41 | 2215.35 | 1.54 |
| ATGC0820 | 1805545 | 1.16 | 5490.10 | 3903.54 | 1.41 |
| ATGC0831 | 2795601 | 1.66 | 14935.85 | 8547.19 | 1.75 |
| ATGC0835 | 2483714 | 1.17 | 2349.16 | 1934.27 | 1.21 |
| ATGC0838 | 4487867 | 1.28 | 48952.51 | 18371.19 | 2.66 |
| ATGC0840 | 1744112 | 0.92 | 5270.01 | 4865.63 | 1.08 |
| ATGC0844 | 1984762 | 1.36 | 6022.43 | 3414.47 | 1.76 |
| ATGC0845 | 3571790 | 1.21 | 8536.37 | 5197.17 | 1.64 |
| ATGC0846 | 6472987 | 0.80 | 40896.39 | 27123.42 | 1.51 |
| ATGC0847 | 5913780 | 2.13 | 5086.51 | 3919.32 | 1.30 |
| ATGC0848 | 1752258 | 0.74 | 8435.69 | 5704.21 | 1.48 |
| ATGC0849 | 6272366 | 1.64 | 15025.88 | 11574.19 | 1.30 |

| | | | | | |
|---|---|---|---|---|---|
| ATGC0851 | 7816689 | 1.14 | 12883.19 | 7882.22 | 1.63 |
| ATGC0852 | 8073621 | 0.94 | 36589.20 | 25607.78 | 1.43 |
| ATGC0853 | 3917759 | 1.07 | 9103.94 | 5019.55 | 1.81 |
| ATGC0855 | 479868 | 0.49 | 589.68 | 477.00 | 1.24 |
| ATGC0857 | 5306277 | 1.19 | 11476.97 | 6263.30 | 1.83 |
| ATGC0858 | 3590716 | 1.66 | 5659.54 | 3131.42 | 1.81 |
| ATGC0859 | 1660951 | 0.73 | 3053.83 | 2184.14 | 1.40 |
| ATGC0860 | 2921657 | 1.57 | 4531.60 | 4537.25 | 1.00 |
| ATGC0861 | 7945061 | 0.82 | 18126.48 | 16907.00 | 1.07 |
| ATGC0862 | 6221248 | 2.21 | 5106.87 | 2636.57 | 1.94 |
| ATGC0863 | 4992805 | 1.69 | 10001.90 | 6131.99 | 1.63 |
| ATGC0864 | 6119030 | 0.97 | 8822.41 | 7210.19 | 1.22 |
| ATGC0865 | 4741350 | 1.20 | 3037.51 | 2013.82 | 1.51 |
| ATGC0866 | 3349943 | 1.49 | 3680.90 | 3092.75 | 1.19 |
| ATGC0867 | 4329931 | 1.34 | 2850.56 | 1653.91 | 1.72 |
| ATGC0868 | 4052258 | 1.67 | 5026.78 | 3115.15 | 1.61 |
| ATGC0869 | 4040012 | 0.73 | 8066.07 | 6679.55 | 1.21 |
| ATGC0871 | 5080660 | 1.13 | 7937.40 | 5527.11 | 1.44 |
| ATGC0872 | 5076637 | 1.34 | 4764.85 | 3707.46 | 1.29 |
| ATGC0873 | 7171767 | 1.41 | 4340.25 | 2783.96 | 1.56 |
| ATGC0874 | 9320089 | 1.02 | 23609.27 | 17885.07 | 1.32 |
| ATGC0875 | 5788809 | 0.81 | 9665.51 | 6040.28 | 1.60 |
| ATGC0876 | 5218828 | 1.78 | 7986.23 | 4436.40 | 1.80 |
| ATGC0877 | 9306871 | 0.66 | 14891.67 | 10419.16 | 1.43 |
| ATGC0878 | 5921869 | 1.93 | 5730.48 | 3398.16 | 1.69 |
| ATGC0879 | 3360630 | 1.41 | 7496.98 | 4679.48 | 1.60 |
| ATGC0880 | 1160484 | 0.88 | 15949.47 | 7746.51 | 2.06 |
| ATGC0881 | 2663111 | 1.12 | 2930.82 | 1940.40 | 1.51 |
| ATGC0882 | 3478815 | 1.16 | 4870.28 | 3230.05 | 1.51 |
| ATGC0883 | 7169141 | 1.11 | 13972.88 | 9132.44 | 1.53 |
| ATGC0884 | 6734763 | 0.76 | 3971.63 | 2911.95 | 1.36 |
| ATGC0885 | 3166259 | 0.91 | 5697.67 | 3547.03 | 1.61 |
| ATGC0886 | 3870385 | 0.72 | 7512.22 | 5101.54 | 1.47 |
| ATGC0887 | 3545181 | 1.64 | 728.05 | 426.38 | 1.71 |
| ATGC0888 | 5366977 | 0.68 | 8200.62 | 5228.78 | 1.57 |
| ATGC0889 | 5260882 | 1.08 | 3554.88 | 2199.48 | 1.62 |
| ATGC0890 | 6421998 | 1.02 | 11658.88 | 7867.20 | 1.48 |
| ATGC0891 | 2854808 | 0.79 | 3940.73 | 2449.27 | 1.61 |
| ATGC0892 | 4507493 | 0.68 | 4919.14 | 3143.28 | 1.56 |
| ATGC0893 | 2415533 | 0.95 | 1611.28 | 946.03 | 1.70 |

| | | | | | |
|---|---|---|---|---|---|
| ATGC0894 | 4961663 | 1.10 | 3453.17 | 2550.81 | 1.35 |
| ATGC0895 | 4264855 | 0.81 | 4238.92 | 2771.78 | 1.53 |
| ATGC0896 | 3271299 | 1.32 | 3110.90 | 2263.43 | 1.37 |
| ATGC0897 | 9899183 | 0.80 | 18829.42 | 14738.07 | 1.28 |
| ATGC0898 | 7772587 | 0.82 | 14228.99 | 9515.81 | 1.50 |
| ATGC0899 | 9621721 | 0.90 | 17999.32 | 12915.67 | 1.39 |
| ATGC0900 | 4519151 | 1.54 | 5901.09 | 4311.01 | 1.37 |
| ATGC0901 | 2391777 | 1.12 | 8466.87 | 5996.34 | 1.41 |
| ATGC0902 | 1375770 | 0.71 | 1537.72 | 1009.73 | 1.52 |
| ATGC0903 | 5838389 | 0.92 | 13784.28 | 8693.62 | 1.59 |
| ATGC0904 | 5693067 | 0.76 | 11697.50 | 7386.13 | 1.58 |
| ATGC0905 | 632370 | 0.31 | 281.95 | 161.72 | 1.74 |
| ATGC0906 | 8033415 | 1.02 | 9831.17 | 7007.56 | 1.40 |
| ATGC0907 | 8112061 | 1.15 | 19248.41 | 15376.84 | 1.25 |
| ATGC0908 | 8188347 | 1.21 | 8824.02 | 6809.78 | 1.30 |
| ATGC0909 | 4786507 | 1.27 | 4676.63 | 4131.57 | 1.13 |
| ATGC0910 | 6240880 | 0.93 | 8783.65 | 5337.13 | 1.65 |
| ATGC0911 | 4171110 | 0.73 | 6631.21 | 3601.44 | 1.84 |
| ATGC0913 | 136065 | 0.30 | 66.46 | 80.86 | 0.82 |
| ATGC0914 | 7965264 | 0.87 | 17854.46 | 12223.40 | 1.46 |
| ATGC0915 | 2504349 | 0.96 | 4608.73 | 3018.79 | 1.53 |
| ATGC0916 | 4009526 | 1.08 | 1895.11 | 953.27 | 1.99 |
| ATGC0917 | 517860 | 0.35 | 188.05 | 100.61 | 1.87 |
| ATGC0918 | 2810923 | 0.90 | 5336.68 | 3102.19 | 1.72 |
| ATGC0919 | 8308430 | 0.85 | 15219.40 | 10511.06 | 1.45 |
| ATGC0920 | 5576178 | 0.62 | 6893.17 | 4510.48 | 1.53 |
| ATGC0940 | 1380398 | 1.89 | 1822.96 | 1403.52 | 1.30 |
| ATGC0944 | 1731110 | 0.82 | 14018.62 | 9092.69 | 1.54 |
| ATGC0946 | 2739240 | 1.36 | 11544.88 | 6756.96 | 1.71 |
| ATGC0948 | 9929346 | 1.32 | 60567.99 | 43586.11 | 1.39 |
| ATGC0952 | 2158673 | 0.92 | 3348.08 | 1899.61 | 1.76 |
| ATGC0953 | 2192574 | 0.86 | 13403.53 | 7834.05 | 1.71 |
| ATGC0954 | 3446446 | 1.61 | 5484.70 | 4287.39 | 1.28 |
| ATGC0955 | 9734476 | 0.83 | 49763.22 | 30207.71 | 1.65 |
| ATGC0956 | 4423100 | 1.47 | 10982.99 | 7257.81 | 1.51 |
| ATGC0957 | 4766714 | 1.33 | 10556.35 | 7709.45 | 1.37 |
| ATGC0958 | 3784450 | 0.85 | 15841.79 | 10807.39 | 1.47 |
| ATGC0959 | 5781564 | 1.58 | 15882.42 | 12305.64 | 1.29 |
| ATGC0960 | 4747744 | 1.14 | 16041.35 | 11206.00 | 1.43 |
| ATGC0961 | 756527 | 0.28 | 7063.01 | 4531.86 | 1.56 |

| | | | | | |
|---|---|---|---|---|---|
| ATGC0962 | 2366130 | 0.81 | 8607.17 | 4674.11 | 1.84 |
| ATGC0964 | 2691371 | 1.34 | 4137.25 | 3033.25 | 1.36 |
| ATGC0965 | 5493098 | 1.75 | 18346.93 | 13088.76 | 1.40 |
| ATGC0966 | 2500544 | 0.97 | 4995.51 | 3023.97 | 1.65 |
| ATGC0967 | 1854966 | 1.20 | 5321.04 | 2595.59 | 2.05 |
| ATGC0968 | 1792919 | 1.23 | 3153.45 | 1964.06 | 1.61 |
| ATGC0969 | 7062790 | 1.66 | 10160.35 | 6346.57 | 1.60 |
| ATGC0970 | 7456404 | 1.07 | 8364.13 | 5715.09 | 1.46 |
| ATGC0971 | 3385517 | 1.01 | 10393.12 | 6362.14 | 1.63 |
| ATGC0972 | 8181934 | 0.82 | 34878.06 | 21931.53 | 1.59 |
| ATGC0973 | 8012773 | 0.97 | 18256.25 | 10729.74 | 1.70 |
| ATGC0974 | 4787829 | 0.80 | 10985.88 | 6612.09 | 1.66 |
| ATGC0975 | 5899149 | 0.94 | 12831.05 | 8660.19 | 1.48 |
| ATGC0976 | 4090554 | 0.85 | 19092.68 | 11777.24 | 1.62 |
| ATGC0977 | 3479000 | 1.13 | 5239.99 | 3178.06 | 1.65 |
| ATGC0978 | 174016 | 0.42 | 174.12 | 95.38 | 1.83 |
| ATGC0979 | 10164010 | 0.97 | 42434.86 | 26434.87 | 1.61 |
| ATGC0980 | 3631970 | 1.27 | 4745.50 | 2944.46 | 1.61 |
| ATGC0981 | 3247380 | 1.24 | 2278.94 | 1512.91 | 1.51 |
| ATGC0982 | 3972953 | 0.88 | 9629.18 | 5918.49 | 1.63 |
| ATGC0984 | 1615932 | 1.69 | 1052.82 | 637.94 | 1.65 |
| ATGC0985 | 5312864 | 0.98 | 36040.06 | 25538.50 | 1.41 |
| ATGC0986 | 3442520 | 0.97 | 10616.49 | 4383.37 | 2.42 |
| ATGC0987 | 648141 | 0.49 | 431.45 | 319.21 | 1.35 |
| ATGC0988 | 3455784 | 1.45 | 5876.45 | 3421.32 | 1.72 |
| ATGC0989 | 3736022 | 1.44 | 2132.79 | 1259.04 | 1.69 |
| ATGC0990 | 8222268 | 0.92 | 39862.77 | 30465.45 | 1.31 |
| ATGC0991 | 4891382 | 0.87 | 10397.10 | 6238.93 | 1.67 |
| ATGC0992 | 4028836 | 1.37 | 4221.24 | 2399.63 | 1.76 |
| ATGC0993 | 7664578 | 1.26 | 28366.46 | 19276.91 | 1.47 |
| ATGC0994 | 820033 | 0.57 | 252.40 | 163.94 | 1.54 |
| ATGC0995 | 465038 | 0.39 | 138.64 | 78.03 | 1.78 |
| ATGC0996 | 4803625 | 1.62 | 11403.73 | 6624.10 | 1.72 |
| ATGC0997 | 5457572 | 1.06 | 6044.76 | 3798.82 | 1.59 |
| ATGC0998 | 3749545 | 0.80 | 3245.41 | 2002.81 | 1.62 |
| ATGC0999 | 4320269 | 1.88 | 5813.48 | 2854.88 | 2.04 |
| ATGC1000 | 4363290 | 1.72 | 4913.65 | 3425.90 | 1.43 |
| ATGC1001 | 643968 | 0.36 | 157.37 | 143.74 | 1.09 |
| ATGC1002 | 5206770 | 1.69 | 3598.99 | 2383.66 | 1.51 |
| ATGC1003 | 4293232 | 1.67 | 10344.46 | 5400.93 | 1.92 |

| | | | | | |
|---|---|---|---|---|---|
| ATGC1004 | 7520872 | 1.13 | 21665.65 | 12133.54 | 1.79 |
| ATGC1005 | 4570058 | 1.35 | 8129.62 | 4673.26 | 1.74 |
| ATGC1006 | 5836284 | 1.65 | 1826.49 | 1259.10 | 1.45 |
| ATGC1007 | 4376282 | 1.50 | 3445.82 | 2223.54 | 1.55 |
| ATGC1008 | 2653044 | 0.74 | 1870.26 | 1117.59 | 1.67 |
| ATGC1009 | 7174823 | 0.97 | 11639.12 | 8267.60 | 1.41 |
| ATGC1010 | 2636834 | 0.98 | 2490.44 | 1436.76 | 1.73 |
| ATGC1011 | 3887657 | 1.27 | 4980.44 | 2777.74 | 1.79 |
| ATGC1012 | 4316928 | 1.40 | 5990.67 | 3844.93 | 1.56 |
| ATGC1013 | 7451216 | 1.04 | 27178.84 | 16615.20 | 1.64 |
| ATGC1014 | 6620423 | 0.86 | 12914.74 | 8757.48 | 1.47 |
| ATGC1015 | 6006526 | 1.31 | 5517.67 | 3806.75 | 1.45 |
| ATGC1016 | 4431096 | 1.22 | 3564.83 | 2202.40 | 1.62 |
| ATGC1017 | 4147522 | 1.23 | 6183.59 | 3293.27 | 1.88 |
| ATGC1018 | 9974036 | 1.00 | 7335.16 | 4168.25 | 1.76 |
| ATGC1019 | 4263354 | 0.95 | 7197.73 | 5327.29 | 1.35 |
| ATGC1020 | 7387177 | 0.80 | 10090.08 | 6466.58 | 1.56 |
| ATGC1021 | 3278404 | 1.31 | 2251.90 | 1434.02 | 1.57 |
| ATGC1022 | 6863786 | 1.00 | 6129.42 | 3751.96 | 1.63 |
| ATGC1023 | 7790908 | 0.91 | 1992.65 | 1404.78 | 1.42 |
| ATGC1024 | 4529080 | 0.89 | 8716.77 | 4970.59 | 1.75 |
| ATGC1025 | 6187229 | 0.95 | 4123.47 | 2637.61 | 1.56 |
| ATGC1026 | 5134005 | 0.81 | 20560.60 | 13179.99 | 1.56 |
| ATGC1027 | 4761506 | 1.18 | 12127.93 | 6522.41 | 1.86 |
| ATGC1028 | 5063997 | 1.77 | 7443.67 | 4293.04 | 1.73 |
| ATGC1029 | 3276692 | 0.98 | 5042.25 | 3277.16 | 1.54 |
| ATGC1030 | 3845378 | 1.00 | 7187.85 | 4886.55 | 1.47 |
| ATGC1031 | 9887354 | 0.85 | 16947.39 | 11117.21 | 1.52 |
| ATGC1032 | 4988435 | 1.24 | 4339.26 | 2574.29 | 1.69 |
| ATGC1033 | 5740952 | 1.18 | 5169.64 | 3155.48 | 1.64 |
| ATGC1034 | 3402498 | 1.44 | 3183.09 | 1770.85 | 1.80 |
| ATGC1035 | 3680998 | 0.98 | 5727.45 | 4047.03 | 1.42 |
| ATGC1036 | 6166834 | 0.72 | 12034.63 | 7535.89 | 1.60 |
| ATGC1037 | 4817150 | 0.88 | 11044.24 | 6644.15 | 1.66 |
| ATGC1038 | 4774789 | 1.02 | 5222.81 | 2619.69 | 1.99 |
| ATGC1039 | 5113117 | 1.81 | 1988.07 | 1276.86 | 1.56 |
| ATGC1040 | 4927880 | 1.58 | 2492.79 | 1757.22 | 1.42 |
| ATGC1041 | 2331134 | 1.18 | 3401.78 | 1916.49 | 1.78 |
| ATGC1042 | 2241290 | 0.56 | 3772.04 | 2147.90 | 1.76 |
| ATGC1043 | 4056935 | 1.47 | 3332.50 | 1632.35 | 2.04 |

| | | | | | |
|---|---|---|---|---|---|
| ATGC1044 | 3440210 | 0.92 | 7665.66 | 4052.76 | 1.89 |
| ATGC1045 | 111994 | 0.62 | 168.22 | 115.65 | 1.45 |
| ATGC1046 | 1433312 | 1.37 | 845.31 | 533.08 | 1.59 |
| ATGC1047 | 3469576 | 1.25 | 2158.67 | 1514.65 | 1.43 |
| ATGC1048 | 5546236 | 0.80 | 3316.08 | 2207.98 | 1.50 |
| ATGC1049 | 4048450 | 1.42 | 5037.13 | 3912.80 | 1.29 |
| ATGC1050 | 4971048 | 1.01 | 6104.35 | 3645.05 | 1.67 |
| ATGC1051 | 4377195 | 1.61 | 9034.64 | 6138.59 | 1.47 |
| ATGC1052 | 3902732 | 1.37 | 2555.15 | 1759.47 | 1.45 |
| ATGC1053 | 2619513 | 0.66 | 2700.74 | 1526.50 | 1.77 |
| ATGC1054 | 4428978 | 0.50 | 4070.29 | 2608.03 | 1.56 |
| ATGC1055 | 4587509 | 1.28 | 6558.45 | 4119.27 | 1.59 |
| ATGC1056 | 4602856 | 1.02 | 8484.98 | 5089.37 | 1.67 |
| ATGC1057 | 3659726 | 0.70 | 4491.31 | 2899.71 | 1.55 |
| ATGC1058 | 1247771 | 0.20 | 4752.78 | 921.10 | 5.16 |
| ATGC1059 | 1099220 | 3.12 | 1315.18 | 797.94 | 1.65 |
| ATGC1060 | 3515026 | 0.88 | 9969.16 | 6336.98 | 1.57 |
| ATGC1061 | 3073103 | 0.73 | 5017.90 | 3073.87 | 1.63 |
| ATGC1062 | 6186282 | 1.20 | 6580.99 | 3785.90 | 1.74 |
| ATGC1063 | 5082926 | 1.08 | 8966.27 | 5630.85 | 1.59 |
| ATGC1064 | 2907314 | 0.38 | 2703.97 | 1212.73 | 2.23 |
| ATGC1065 | 2755522 | 1.26 | 4570.81 | 2902.63 | 1.57 |
| ATGC1066 | 6453736 | 0.99 | 8803.44 | 5617.45 | 1.57 |
| ATGC1067 | 5188586 | 0.71 | 2726.62 | 1680.40 | 1.62 |
| ATGC1068 | 653158 | 0.42 | 268.46 | 210.74 | 1.27 |
| ATGC1069 | 5014828 | 0.98 | 1679.62 | 1020.80 | 1.65 |
| ATGC1070 | 2901872 | 1.59 | 4328.29 | 2863.30 | 1.51 |
| ATGC1071 | 9687506 | 0.87 | 29406.49 | 21031.98 | 1.40 |
| ATGC1072 | 3409622 | 0.94 | 2707.54 | 1800.95 | 1.50 |
| ATGC1073 | 4310163 | 0.88 | 6243.38 | 3586.10 | 1.74 |
| ATGC1074 | 3852812 | 0.82 | 9008.00 | 5176.60 | 1.74 |
| ATGC1075 | 3635750 | 0.87 | 2270.92 | 1758.24 | 1.29 |
| ATGC1078 | 1642612 | 1.42 | 3225.23 | 1759.39 | 1.83 |
| ATGC1079 | 2121360 | 1.31 | 4528.41 | 2616.71 | 1.73 |
| ATGC1080 | 2525132 | 0.97 | 10676.35 | 5853.50 | 1.82 |
| ATGC1081 | 4806776 | 0.72 | 8748.85 | 5680.72 | 1.54 |

**Table S2. Estimates of selective constraints in 30 splittable ATGCs.**

| ATGC ID | Constraint - Subtree 1 | Constraint - Subtree 2 | Ratio |
|---|---|---|---|
| ATGC0056 | 1.37 | 0.47 | 2.94 |
| ATGC0008 | 1.13 | 0.39 | 2.91 |
| ATGC0053 | 1.63 | 0.96 | 1.71 |
| ATGC0284 | 1.56 | 1.02 | 1.52 |
| ATGC0134 | 1.42 | 0.98 | 1.45 |
| ATGC0460 | 1.29 | 0.89 | 1.45 |
| ATGC0296 | 0.95 | 0.68 | 1.39 |
| ATGC0462 | 1.18 | 0.88 | 1.34 |
| ATGC0004 | 0.82 | 0.65 | 1.27 |
| ATGC0111 | 1.78 | 1.42 | 1.25 |
| ATGC0149 | 1.48 | 1.18 | 1.25 |
| ATGC0024 | 0.75 | 0.61 | 1.23 |
| ATGC0184 | 0.74 | 0.61 | 1.22 |
| ATGC0143 | 0.71 | 0.61 | 1.17 |
| ATGC0265 | 1.16 | 1.00 | 1.17 |
| ATGC0104 | 0.62 | 0.53 | 1.15 |
| ATGC0144 | 0.50 | 0.44 | 1.14 |
| ATGC0089 | 1.03 | 0.91 | 1.13 |
| ATGC0402 | 1.22 | 1.10 | 1.10 |
| ATGC0403 | 0.70 | 0.64 | 1.09 |
| ATGC0059 | 1.28 | 1.17 | 1.09 |
| ATGC0001 | 1.18 | 1.08 | 1.09 |
| ATGC0071 | 1.23 | 1.14 | 1.07 |
| ATGC0015 | 0.82 | 0.76 | 1.07 |
| ATGC0116 | 1.01 | 0.94 | 1.07 |
| ATGC0290 | 1.04 | 0.97 | 1.07 |
| ATGC0002 | 1.25 | 1.18 | 1.06 |
| ATGC0100 | 1.28 | 1.26 | 1.01 |
| ATGC0098 | 1.44 | 1.42 | 1.01 |
| ATGC0124 | 0.87 | 0.86 | 1.00 |

**Table S3. Correlation between genome size and ATGC-specific selective constraint across various prokaryotic clades. Only clades consisting of more than 15 genomes were included**

| Taxon | Count | Spearman's ρ | CI lower | CI upper | p_value |
|---|---|---|---|---|---|
| Bacteria | 772 | 0.28 | 0.20 | 0.35 | 6.07E-15 |
| Archaea | 19 | -0.82 | -0.93 | -0.64 | 1.78E-05 |
| *Pseudomonadota* | 323 | 0.38 | 0.27 | 0.49 | 1.10E-12 |
| *Actinomycetota* | 152 | -0.09 | -0.26 | 0.09 | 2.54E-01 |
| *Bacillota* | 144 | 0.00 | -0.15 | 0.17 | 9.86E-01 |
| *Bacteroidota* | 60 | 0.35 | 0.09 | 0.59 | 5.60E-03 |
| *Campylobacterota* | 20 | 0.21 | -0.35 | 0.61 | 3.80E-01 |
| *Cyanobacteriota* | 19 | -0.17 | -0.55 | 0.28 | 4.82E-01 |
| *Mycoplasmatota* | 17 | 0.07 | -0.44 | 0.60 | 7.79E-01 |
| *Gammaproteobacteria* | 172 | 0.53 | 0.39 | 0.65 | 7.10E-14 |
| *Actinomycetes* | 149 | -0.07 | -0.24 | 0.11 | 3.99E-01 |
| *Bacilli* | 109 | 0.12 | -0.07 | 0.31 | 2.12E-01 |
| *Alphaproteobacteria* | 92 | 0.30 | 0.09 | 0.51 | 3.43E-03 |
| *Betaproteobacteria* | 58 | 0.32 | 0.01 | 0.55 | 1.55E-02 |
| *Flavobacteriia* | 36 | 0.58 | 0.26 | 0.80 | 2.00E-04 |
| *Clostridia* | 26 | -0.01 | -0.40 | 0.42 | 9.78E-01 |
| *Bacillales* | 61 | -0.33 | -0.52 | -0.12 | 9.35E-03 |
| *Enterobacterales* | 58 | 0.48 | 0.19 | 0.69 | 1.38E-04 |
| *Lactobacillales* | 48 | 0.31 | -0.01 | 0.56 | 3.13E-02 |
| *Hyphomicrobiales* | 45 | 0.06 | -0.27 | 0.40 | 7.03E-01 |
| *Mycobacteriales* | 43 | 0.41 | 0.09 | 0.66 | 7.01E-03 |
| *Micrococcales* | 42 | 0.04 | -0.31 | 0.39 | 7.95E-01 |
| *Burkholderiales* | 41 | -0.13 | -0.47 | 0.22 | 4.04E-01 |
| *Kitasatosporales* | 37 | -0.27 | -0.55 | 0.08 | 1.04E-01 |
| *Eubacteriales* | 23 | -0.19 | -0.57 | 0.26 | 3.73E-01 |
| *Pseudomonadales* | 23 | 0.00 | -0.40 | 0.42 | 9.89E-01 |
| *Alteromonadales* | 18 | 0.12 | -0.55 | 0.64 | 6.39E-01 |
| *Vibrionales* | 17 | 0.32 | -0.24 | 0.75 | 2.13E-01 |
| *Lactobacillaceae* | 28 | 0.38 | -0.02 | 0.67 | 4.79E-02 |
| *Bacillaceae* | 23 | 0.19 | -0.27 | 0.54 | 3.83E-01 |
| *Microbacteriaceae* | 22 | -0.05 | -0.51 | 0.47 | 8.28E-01 |
| *Enterobacteriaceae* | 21 | 0.50 | 0.00 | 0.75 | 1.99E-02 |
| *Flavobacteriaceae* | 21 | 0.29 | -0.18 | 0.64 | 2.07E-01 |
| *Pseudomonadaceae* | 19 | 0.14 | -0.30 | 0.56 | 5.81E-01 |
| *Burkholderiaceae* | 17 | -0.21 | -0.67 | 0.42 | 4.11E-01 |
| *Mycobacteriaceae* | 16 | 0.56 | 0.01 | 0.87 | 2.54E-02 |
